# Engineering Basal Cognition: Minimal Genetic Circuits for Habituation, Sensitization, and Massed–Spaced Learning

**DOI:** 10.1101/2025.09.09.674925

**Authors:** Jordi Pla-Mauri, Ricard Solé

## Abstract

Cognition is often associated with complex brains, yet many forms of learning—such as habituation, sensitization, and even spacing effects—have been observed in single cells and aneural organisms. These simple cognitive abilities, despite their cost, offer evolutionary advantages by allowing organisms to reduce environmental uncertainty and improve survival. Recent studies have confirmed early claims of learning-like behavior in protists and slime molds, pointing to the presence of basal cognitive functions long before the emergence of nervous systems. In this work, we adopt a synthetic biology approach to explore how minimal genetic circuits can implement non-associative learning in unicellular and multicellular systems. Building on theoretical models and using well-characterized regulatory elements, we design and simulate synthetic circuits capable of reproducing habituation, sensitization, and the massed–spaced learning effect. Our designs incorporate activators, repressors, fluorescent reporters, and quorum-sensing molecules, offering a platform for experimental validation. By examining the structural and dynamical constraints of these circuits, we highlight the distinct temporal dynamics of gene-based learning systems compared to neural counterparts and provide insights into the evolutionary and engineering challenges of building synthetic cognitive behavior at the cellular level.

## I. INTRODUCTION

Life on Earth has evolved in multiple directions, with major evolutionary events defining the rise of novelties, such as the transition from unicellular to multicellular or the emergence of language (Szathmáry and Smith, 1995; Lane, 2010). A general trait shared by most of these transitions is the emergence of new types of agents capable of dealing with environmental uncertainties in novel ways. Among others, the evolution of neural agents represents a revolutionary path toward neural networks and brains (Ginsburg and Jablonka, 2010). The rise of cognitive structures and brains largely begins with the Cambrian explosion event, where we can see the rapid evolution of animals equipped with mobile parts and sensors (Hsieh *et al*., 2022). Using a special class of cells, the neuron, it was possible to develop mechanisms to store and process information reliably (Monk and Paulin, 2014). In this context, learning might have played a crucial role in the development of complex brains (Ginsburg and Jablonka, 2021).

Cognitive structures are costly, from sensors and actuators to the whole brain. How can evolution favor their emergence? The answer is that mastering time, i.e., storing past information that can be used to predict the environment, can have a high return. In other words, reducing uncertainty has a high pay-off. Not sur-prisingly, memory is a widespread feature of most multicellular life, and several kinds of learning mechanisms have been described (Gunawardena, 2022). These include two well-known processes, namely habituation and sensitization, which have been known for thousands of years (Blumstein, 2016) as well as associative learning. However, these features are not limited to multicellular systems and were identified by several authors regarding the behavioral responses of individual cells (Dussutour, 2021). This is the case of *Stentor* (Figure 1a), a single-cell protozoan (Binet, 1889; Jennings, 1931) which was shown to exhibit an enhanced response to subsequent milder stimuli, that is, *habituation*. This indicated that it was learning to associate the intense stimulus with a potential danger, resulting in increased sensitivity. Similarly, *Stentor* was also shown to exhibit a reduced response under repeated exposure to the same stimulus, indicating that it was learning to recognize the stimulus as non-threatening or irrelevant. In addition, *sensitization*, when an organism becomes more sensitive to a stimulus after experiencing an intense or aversive event, was also observed.

**FIG. 1.**
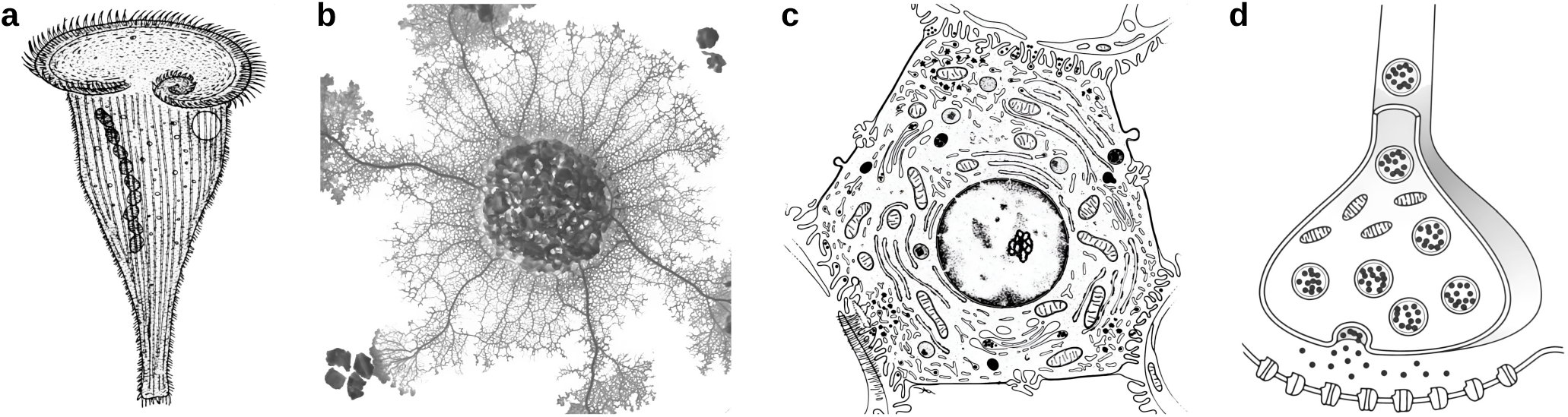
Embodiments for basal cognition. Simple forms of learning (associative and non-associative) in simple biological systems can be observed in free-living unicellular life forms, such as the protozoan *Stentor* (a), the *Physarum polycephalum* slime mold (b), but can also be found in non-neural tissues, such as hepatic cells (c), and has been traditionally reported in neural systems on the small, synaptic scale (d).

Although Jennings work was questioned due to reproducibility issues, recent studies confirmed (and ex-panded) his key results, uncovering a hierarchy of decision-making actions (Eisenstein *et al*., 1982; Dexter *et al*., 2019; Gershman *et al*., 2021; Wright *et al*., 2023).

Both habituation and sensitization, along with other forms of learning, have also been found in *Physarum polycephalum* (Figure 1b). This aneural organism is a giant multinucleated cell that can extend up to hun-dreds of square centimeters. It has been shown to exhibit complex self-organized patterns and solve a wide range of tasks that involve complex computational decisions (Adamatzky, 2010; Bonifaci *et al*., 2012; Adamatzky, 2016; Vallverdú *et al*., 2018).

Basal cognition encompasses the fundamental processes and mechanisms that enable organisms to detect certain environmental states and respond appropriately to ensure survival (such as finding food and avoiding danger) and reproduction long before the evolution of nervous systems. Modern cells have complex molecular networks that can participate in cognitive tasks, from detecting and responding to chemical cues to actively exploring their worlds in space and time. An important aspect of their behavioral repertoires is the presence of memory and learning characteristics. They are in fact strong prerequisites to build a behavior. In recent years, renewed interest has emerged in this area, including dedicated efforts to operationally reframe cognition (Lyon *et al*., 2021).

Theoretical and computational studies on these simple forms of learning have recently been developed, searching for potential circuits capable of implementing them within cells, particularly in terms of habituation (Smart *et al*., 2024; Eckert *et al*., 2024). There is an alternative approach to this problem from an engineering perspective. In general, while much understanding of major evolutionary transitions has been gained through molecular phylogenetics or comparative analysis and paleobiological data, we can also address the problem by recreating these evolutionary events and their precursors using (among other approaches) synthetic biology (Sole, 2016). In this context, the design of genetic circuits and the potential of tissue bioengineering can help understand the constraints associated with biological complexity. This includes morphology (Davies, 2008; Ebrahimkhani and Levin, 2021; Davies and Levin, 2023), multicellu-larity (Maharbiz, 2012; Solé *et al*., 2024), ecology (Mee and Wang, 2012; Solé *et al*., 2024), biorobots (Blackiston *et al*., 2023) or collective intelligence (Solé *et al*., 2016), among other problems.

Previous studies have addressed the problem of classical conditioning (associative learning) circuits based on transcriptional network systems, including several proposals for candidate molecular circuits (Fernando *et al*., 2009; Lu *et al*., 2009; Sorek *et al*., 2013; Macia *et al*., 2017; Biswas *et al*., 2022). In this paper, we take the “synthetic” approach to consider how simple genetic circuits could reliably implement non-associative learning, thus involving a single type of stimulus. This will include the two previous case studies (habituation and sensitization) and the so-called massed–spaced learning effect. The latter has traditionally been discussed within the psychology literature and involves scenarios in which long-term memory is enhanced when learning events are spaced apart in time, rather than occurring in immediate succession. However, recent studies have shown that it can also be present in single human cells (Kukushkin *et al*., 2024), suggesting that this learning effect might also operate on a small scale within whole bodies.

Our implementations rely on well-characterized genetic parts and regulatory mechanisms, making them amenable to experimental realization. The designed circuits proposed here will include repressors, fluorescent reporters, and quorum-sensing molecules as key components, allowing external control of the input signal and measurement of the average output at the population level. By following our synthetic approach, there is a marked difference between the gene networks that implement learning and their synaptic counterparts: a much slower time scale. This difference will be relevant to our discussion on the role that learning plays in single-cell organisms and the constraints imposed on engineering designs.

## II. RESULTS

### A. Habituation

Habituation represents one of the simplest and most ubiquitous forms of non-associative learning, characterized by a gradual decrease in the behavioral or physiological response after repeated exposure to a stimulus that is perceived to be neither harmful nor beneficial (Thompson and Spencer, 1966). This phenomenon is not merely passive fatigue or sensory adaptation, but an active process. It has been extensively studied across various species—from *Aplysia* to humans—demonstrating its evolutionary conservation and functional importance in adaptive behavior (Rankin *et al*., 2009).

This fundamental mechanism of behavioral adaptation allows organisms to efficiently filter out irrelevant or non-threatening stimuli from their environment, such as background noise or persistent visual cues. By reducing responses to familiar inputs, habituation enables the organism to conserve cognitive and energetic resources, which can then be redirected toward detecting and responding to novel or potentially significant environmental changes (Pavlov, 2010). In this way, habituation plays a crucial role in attentional modulation and information processing, helping organisms prioritize survival-relevant stimuli.

At its core, habituation requires a responsive system capable of detecting external stimuli and modulating its sensitivity over time. This modulation typically involves negative feedback mechanisms that accumulate with repeated stimulation. For example, repeated activation of postsynaptic receptors may lead to internalization of these receptors or alterations in second-messenger signaling pathways, effectively raising the threshold required to elicit a response (Kandel, 2001). Such mechanisms enable the system to adaptively ignore persistent signals while remaining sensitive to new or changing inputs.

Figure 2a sketches the logic of our proposed minimal circuit, and in Figure 2b a genetic design is shown as a realization of an incoherent feed-forward loop (I-FFL), a motif previously discussed in the literature (Eckert *et al*., 2024; Staddon, 1993; Staddon and Higa, 1996). Alternative designs based on negative feedback topologies for the receptor are discussed in the Supplementary Material (SM I.B, I.C), where they are shown to perform less optimally.

**FIG. 2.**
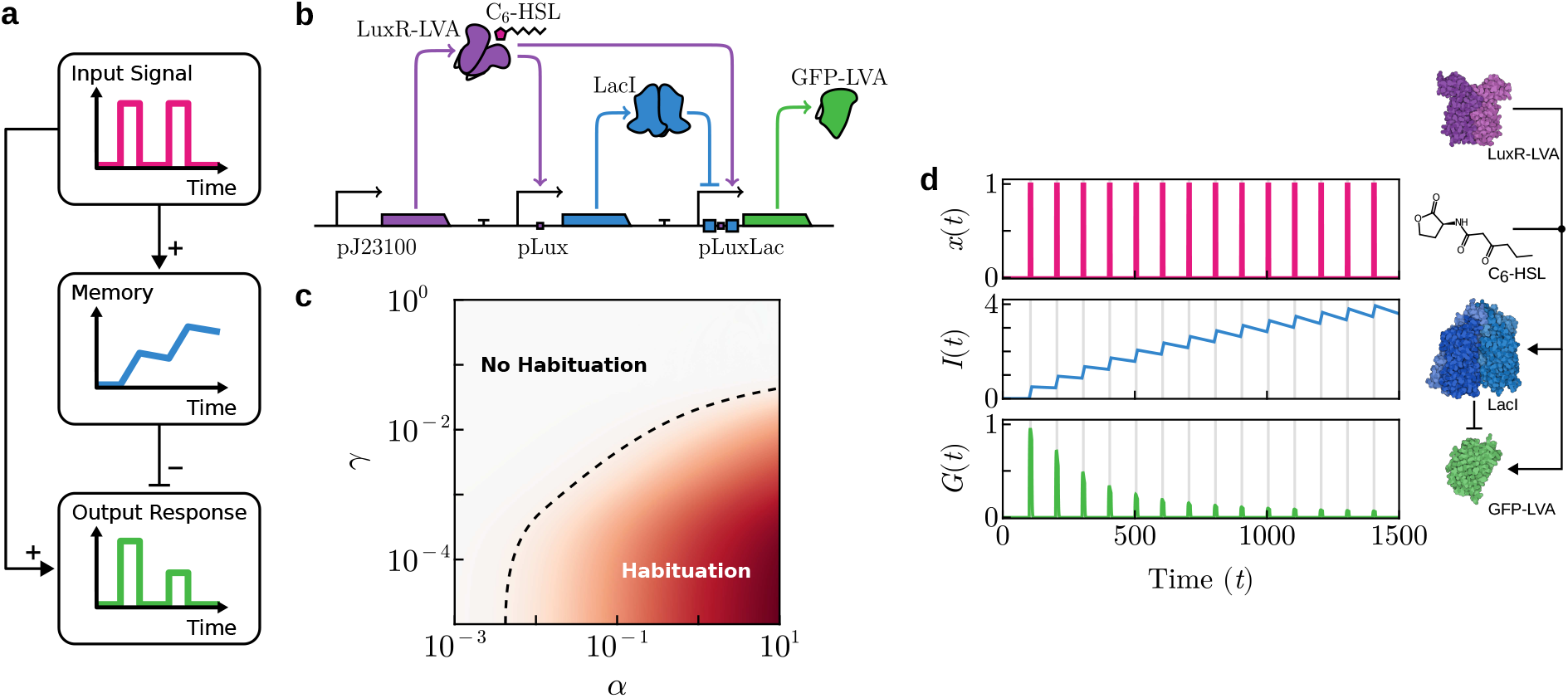
Synthetic circuit design and behavior for habituation in a single-cell. In (a), the basic logic requirements for habituation are outlined in a causal diagram. In (b), an explicit synthetic circuit design is shown. A constitutively expressed LuxR-LVA receptor binds to C_6_-HSL input molecules. This complex activates LacI repressor expression. The output, GFP-LVA, requires both an absence of the repressor and the presence of the input-receptor complex for activation. In (c), a parameter space showing habituation strength, measured as a log_2_-fold change (FC) between the last and the first peak. In (d), a sample time-series simulation showing habituation response following equations (1). Vertical gray regions indicate the presence of an external stimulus *x*. Key molecular components of the circuit are shown on the right, along with their interactions. Protein structures were rendered using Illustrate (Goodsell *et al*., 2019). Parameters: *α* = 10^−1^, *β* = 2, *γ* = 10^−3^, *λ* = 1, *µ* = 1, Δ*τ*_on_ = 10, Δ*τ* = 100.

The proposed synthetic circuit senses and reacts to an external input signal *x*, such as the quorum sensing molecule C_6_-HSL, which diffuses into the cell and binds to a constitutively expressed transcriptional regulator LuxR-LVA, denoted *X*. The resulting ligand-bound complex *X*–*x* activates transcription of two key components: a memory *I* and an output *G*.

The memory component *I*, the repressor protein LacI in the example, is expressed under an inducible promoter activated by *X*–*x*, and its concentration accumulates over successive input pulses, serving as a biochemical integrator of past input activity. Output *G*, represented for simplicity as the fluorescent reporter GFP-LVA, is expressed under a hybrid promoter that requires the presence of complex *X*–*x* for activation while simultaneously being repressed by *I*, which approximates N-IMPLY logic (*x* ∧ ¬*I*). However, rather than enforcing a sharp binary switch, intermediate concentrations of *I* lead to partial suppression of promoter activity, effectively reducing the output production rate.

All proteins except memory *I* contain a degradation tag that enhances their proteolytic turnover, resulting in a higher degradation rate. This difference in stability between sets of proteins establishes a clear separation of time scales, ideally allowing only untagged proteins to persist between input pulses.

The equations governing the system are as follows:

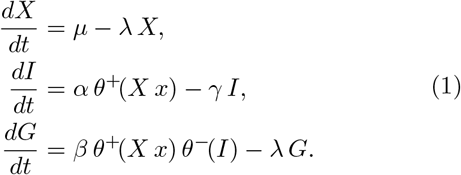

The key parameters include the maximal expression rates *α, β*, and *µ*. These parameters encapsulate the combined strength of a promoter and its RBS, allowing each to be independently tuned. The basal degradation-dilution rate *γ* governs the turnover of untagged proteins, while *λ* denotes the higher degradation rate of tagged proteins.

Functions *θ*^±^(·) are normalized transfer functions of the promoters, defined as standard Hill form with half-activation constant *K*_1*/*2_ = 1, namely:

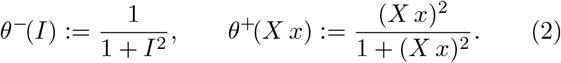

To assess the circuit’s capacity to habituate to a periodic pulsating signal, simulations are conducted starting from a steady state with no external input. A periodic external input *x* is introduced as an instantaneous pulse lasting for a duration Δ*τ*_on_, followed by an input-free interval of duration Δ*τ*_off_, allowing the system to relax. The total period of each cycle is therefore Δ*τ* = Δ*τ*_on_ + Δ*τ*_off_. The trajectory is partitioned into a collection of *n* con-secutive intervals 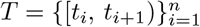, where each *t*_*i*_ cor-responds to the onset of the *i*-th input pulse, and each interval has a constant duration Δ*τ* = *t*_*i*+1_ − *t*_*i*_.

An example of the time-series resulting from this minimal circuit is shown in Figure 2d, where the time-series for variables *x, I* and *G* are displayed. The concentration of repressor *I* shows a growing trend over time, decreasing slightly during relaxation periods as its production halts while continuously decaying with rate *γ*. A habituation pattern is displayed by *G* with regular peaks displaying a decrease in amplitude with each successive input pulse.

The conditions required to guarantee the growth of the memory component can be analytically derived given some approximations. Since a memory component is also relevant for other motifs discussed, a basic derivation is warranted (see also SM II).

To this end, trajectories are partitioned into periodic intervals, each consisting of an active phase of duration Δ*τ*_on_, during which an external input is applied, followed by a relaxation phase of duration Δ*τ*_off_, where no input is present.

Assuming that the memory component does not approach saturation—specifically, that *I*(*t*) ≪ *α/γ* holds throughout the entire trajectory—and further assuming that the duration of each pulse is much shorter than the relaxation period (Δ*τ*_on_ ≪ Δ*τ*_off_), the dynamics of memory accumulation can be effectively approximated as a sequence of instantaneous increments occurring at the onset of each pulse, followed by continuous exponential decay between events.

Under the impulsive approximation, the temporal evolution of the memory component is governed by the following differential equation:

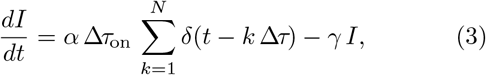

where *δ*(·) denotes the Dirac delta function, representing a sequence of *N* instantaneous inputs applied at discrete times *t* = *k* Δ*τ*.

Assuming negligible initial conditions, memory dynamics are given by:

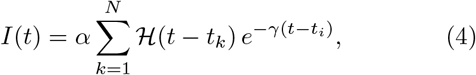

where ℋ (·) denotes the Heaviside step function, a piecewise function defined as 0 for negative arguments and 1 otherwise, and *t*_*k*_ = *k* Δ*τ* are the times where input pulses are applied.

The change in memory over a single interval can be expressed as:

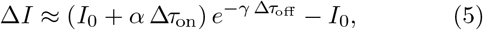

where *I*_0_ represents the initial memory value just before the input is applied.

Since the net change in memory per cycle, Δ*I*, decreases monotonically as the relaxation duration Δ*τ*_off_ increases, and since Δ*I >* 0 when Δ*τ*_off_ = 0 (memory accumulates in the absence of extended downtime), it follows that there exists a critical threshold for Δ*τ*_off_ beyond which no net memory gain occurs over successive stimulation cycles. Therefore, memory accumulation is possible only if the relaxation period satisfies the inequality:

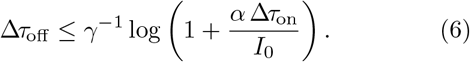

As the number of events increases, *N* → ∞, the periodic steady state solution approximates:

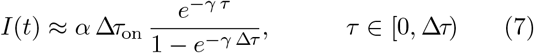

oscillating with period Δ*τ* ≈ Δ*τ*_off_.

In Figure 2c a two-dimensional parameter space (*γ, α*), showing the adaptation strength, quantified as a fold change, FC, defined as the log_2_ ratio between the maximum response after the final stimulus and the peak response observed throughout the simulation time span, which in this case occurs within the first stimulus interval:

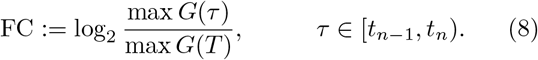

Two domains are clearly defined, with a boundary separating a phase where habituation occurs from another where habituation is not possible.

Robust habituation depends on memory *I* being able to hold its state between stimuli, which can achieved by ensuring it degrades slowly (low *γ*). At the same time, increasing the memory production rate *α* makes habituation stronger by causing the system’s final response to be a smaller fraction of its initial output. However, a higher production rate also reduces the system’s peak response, creating a key trade-off: a stronger habituation strength comes at the cost of a lower maximum output. A more in depth analysis of this trade-off is provided in SM I.A.

### B. Sensitization

Sensitization—also called facilitation—is another fundamental form of non-associative learning that stands in direct contrast to habituation. While habituation involves a decrease in response after repeated exposure to a stimulus, sensitization is characterized by an amplification of the response after repeated or intense stimulation (Kandel, 2001).

This heightened responsiveness serves as an essential adaptive mechanism across a wide range of species, enabling organisms to prioritize and react more vigorously to potentially harmful or biologically significant stimuli (Pinsker *et al*., 1973). Sensitization has been extensively studied in the context of defensive behaviors and nociceptive responses, where it enhances survival by promoting rapid responses to environmental threats. Beyond behavioral responses, sensitization manifests itself at multiple levels of biological organization, from whole-organism reflexes to synaptic plasticity and cellular signaling pathways (Carew *et al*., 1981).

The molecular mechanisms that underlie sensitization are generally based on positive feedback loops or cascade amplification systems, which progressively strengthen in response to repeated stimulation. In this work, a simple synthetic design that recapitulates this behavior is presented, drawing upon elements from our earlier design for habituation. The key to this system lies again in leveraging the molecular memory provided by repressor I, which generates a signal that accumulates in the presence of input and decays slowly during the relaxation phase; this dynamic enables the circuit to effectively “remember” prior stimulations and respond more strongly upon repeated exposure. It is also worth noting that, in the context of synaptic plasticity, sensitization is often achieved through a basic motif derived from the habituation mechanism, with the addition of an extra interneuron (LeDoux, 2003).

Building on the basic logic of sensitization illustrated in Figure 3a, a functional genetic circuit can be derived. This proposed circuit (Figure 3b) largely reuses the gene regulatory interactions from the previous habituation design but incorporates a few key modifications to establish a positive feedback loop.

**FIG. 3.**
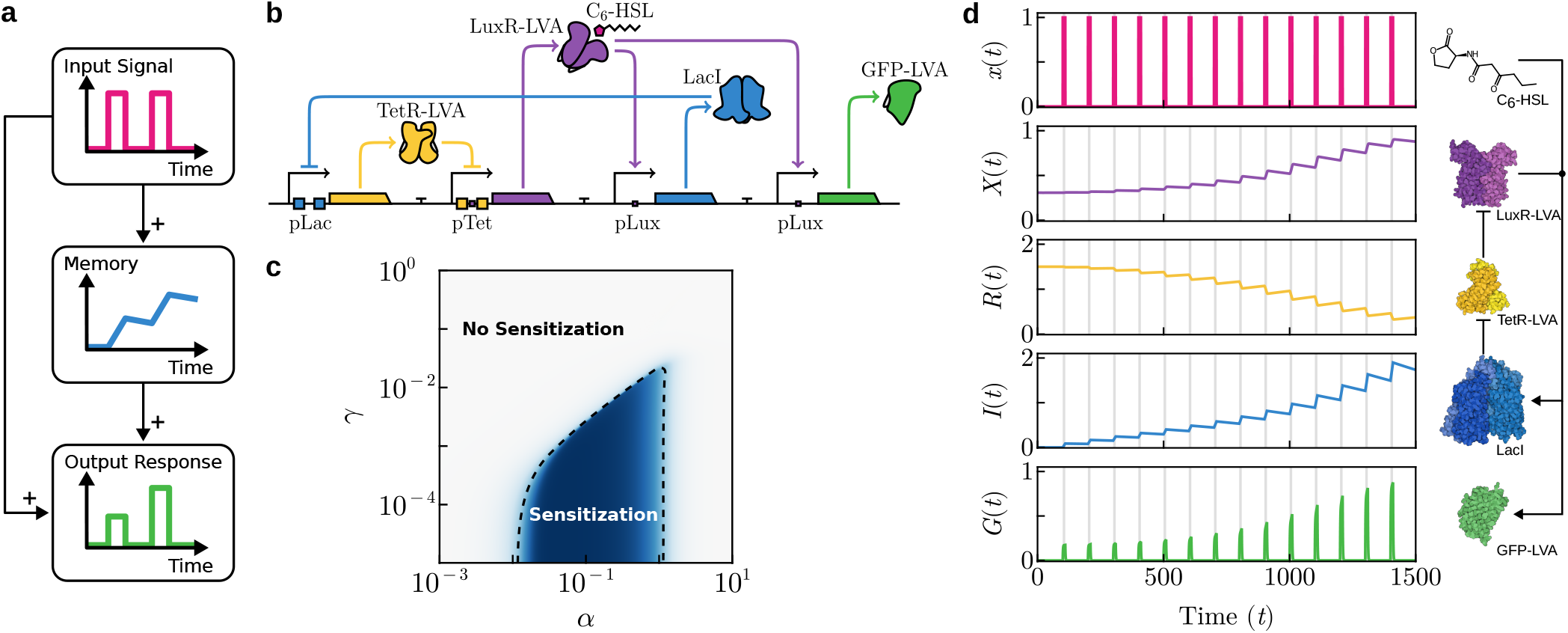
Synthetic circuit design and behavior for sensitization in a single-cell. In (a), the basic logic requirements for sensitization are outlined in a causal diagram. In (b), an explicit synthetic circuit design is shown. When the input molecule (C_6_-HSL) is present, it binds to its receptor (LuxR-LVA), which then simultaneously activates both the repressor (LacI) and the circuit’s output signal (GFP-LVA). Repressor TetR-LVA, which controls LuxR-LVA expression, is itself repressed by LacI, creating an indirect positive regulatory feedback loop that enables the sensitization behavior. In (c), a parameter space showing habituation strength, measured as a log_2_-fold change (FC) between the last and the first peak. In (d), a sample time-series simulation showing sensitization response following equations (9). Vertical gray regions indicate the presence of an external stimulus *x*. Key molecular components of the circuit are shown on the right, along with their interactions. Parameters: *α* = 10^−1^, *β* = 2, *γ* = 10^−3^, *λ* = 1, *µ* = 1, Δ*τ*_on_ = 10, Δ*τ* = 100.

The circuit creates a positive feedback loop on receptor *X* (LuxR-LVA) through a double repression cascade. In the presence of input *x*, the *X*–*x* complex drives expression of repressor *I* (LacI), which in turn represses expression of *R* (TetR-LVA). Since *R* itself represses the production of *X*, the net effect is that *X* indirectly promotes its own expression and amplification. This design decouples the feedback strength through intermediate regulators, allowing tunable and graded amplification that depends on the stimulation history. By using this indirect mechanism, the system avoids some limitations inherent to a direct self-activation loop (see SM III.B).

The output *G* (GFP-LVA), which is expressed from a promoter directly activated by the *X*–*x* complex, is consequently also amplified. Thus, the production of *G* is dependent upon both the presence of the external input and the time-dependent concentration of *X*, ensuring input-gated activation and a sensitized response that reflects the system’s history of stimulation. An alternative design based on a coherent feed-forward loop (C-FFL), which offers similar performance and complexity, is discussed in the Supplementary Material (SM III.C).

The key to the memory effect is that only the repressor *I* has a slow degradation rate (*γ*). Its continued presence represses *R*, which in turn relieves repression of the receptor *X* promoter. The fast degradation rate (*λ*) of both *X* and *R* ensures their rapid turnover, with their concentrations rapidly approaching a quasi-steady-state set by the slower dynamics of *I*, enabling time-scale separation and effective memory encoding.

The equations governing the system are as follows:

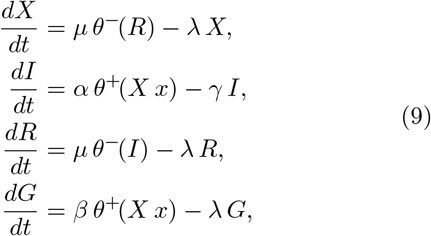

where *α, β, µ* represent independently tunable maximal expression rates, *γ* is the basal rate at which proteins degrade, and *λ* is the degradation rate of short-lived (tagged) proteins. As before, the constraint *λ* ≫ *γ* is assumed, creating a separation of timescales. This ensures that untagged proteins degrade rapidly, while other proteins accumulate within the cell over successive pulses.

An example of a time-series that displays sensitization is shown in Figure 3d. As in the habituation example, the memory component *I* accumulates over time—but here, this leads to a decrease in repressor *R*, which otherwise inhibits production of *X*, effectively boosting receptor levels receptor levels in proportion. This double inhibition loop effectively increases sensitivity to input *x*, yielding progressively larger peaks in output *G*.

Figure 3c shows the parameter space (*α, γ*) for the sensitization model, where the fold change from equation (8) is used to display a well-defined boundary of sensitization responses (blue domain). The sharp decrease in the fold change as the production rate of the memory molecule (*α*) increases is due to the rapid accumulation of memory *I*, which quickly saturates its corresponding transfer function. When *α* is high, the system approaches this saturation limit during the first peak response, thereby limiting any further increase in the levels of the input receptor and the output protein (see SM III.A for details).

### C. Combining Sensitization and Habituation

Cognition is not limited to specific kinds of response, and a relevant question is how can we combine or modify our minimal motifs in such a way that multiple learning responses can be at work. This occurs with some simple neural circuits that can encode both increased and decreased responsiveness. This is the case of classic experiments in *Aplysia californica*, where a slight touch to the siphon triggers a gill-withdrawal reflex (Biswas *et al*., 2022; Rankin and Carew, 1987). After a strong stimulus such as a tail shock, this response becomes sensitized, with even light touches causing a stronger reaction. However, if light touch is repeated without reinforcement, the reflex gradually habituates, weakening over time. While sensitization involves enhanced neurotransmitter release, habituation results from reduced synaptic activity.

In this section, we briefly illustrate the possibility of combining sensitization and habituation responses within a single circuit to produce a hybrid response. This results in an output where an initial period of sensitization is followed by habituation.

Although one might expect this construct to require some nontrivial combination of the previous motifs, surprisingly this more complex behavior does not require a fundamentally new design (Figure 4a). It can be achieved simply by reintroducing a single repressor interaction from the habituation circuit, the hybrid promoter from Figure 2b, into the sensitization design from Figure 3b. In this hybrid circuit, output *G* (GFP-LVA) expression depends on both the presence of the input-bound receptor *X*–*x* and the absence of repressor *I*. The behavioral transition occurs when the inhibitory effect of *I* on output expression dominates over the activation pathway mediated by *X*–*x*.

**FIG. 4.**
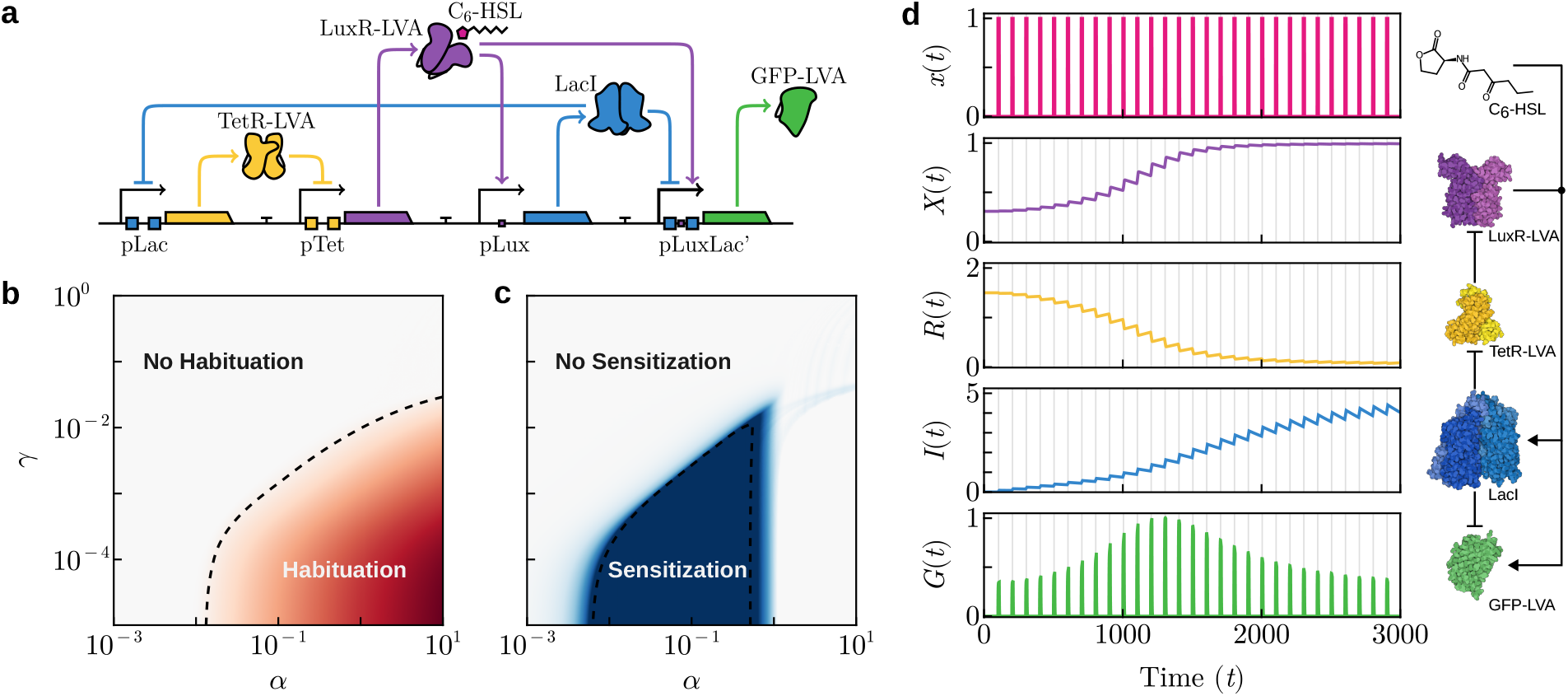
Synthetic circuit design and behavior for sensitization-habituation in a single-cell. In (a), an explicit synthetic circuit design is shown. In (b, c), the corresponding parameter spaces for the habituation and sensitization regimes are shown, with a dashed lane dividing spaces where | FC | *>* 1. The same indicator (FC) is used, but in this case, the first peak is compared to the highest peak to measure sensitization, and the last peak is compared to the highest peak to measure habituation. In (d) the time-series for the model is shown, with sensitization followed by habituation, following equations (10). The vertical gray shading denotes temporal intervals during which external stimulus *x* is present. Key molecular components of the circuit are shown on the right, along with their interactions. Parameters: *α* = 10^−1^, *β* = 4, *γ* = 10^−3^, *λ* = 1, *µ* = 1, Δ*τ*_on_ = 10, Δ*τ* = 100.

The number of pulses required to transition from sensitization to habituation is tuned by the relative sensitivity to *I* for the two promoters regulated by it. If the hybrid promoter requires a higher repressor concentration, a clearer separation between the two phases can be observed. This sensitivity can be experimentally engineered through mutagenesis or predicted *in silico*.

An alternative implementation tweaking a C-FFL sensitization motif is also possible by using two distinct sources for repressor *I*, with different dynamics: a tagged version, indirectly repressed by the input complex for sensitization; and an untagged version, directly expressed by the same complex for habituation (see SM IV.C).

The equations governing the system are as follows:

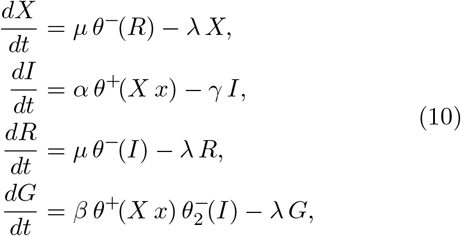

where *α, β, µ* represent independently tunable maximal expression rates, *γ* is the basal rate at which proteins degrade, and *λ* is the degradation rate of short-lived (tagged) proteins.

The function 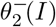 represents the repressor curve for the output. It is analogous to the repressor transfer function presented in Equation (2), but with a doubled half-activation constant:

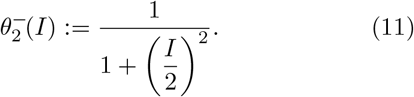

This increase in *K*_1*/*2_ shifts the system’s dynamic response to a longer characteristic time scale. Consequently, the point at which the system’s behavior shifts from sensitization to habituation occurs more slowly.

Figure 4d illustrates an example time-series in which the memory component *I* accumulates progressively over time, as observed in earlier cases. As *I* builds up, receptor *X* increases, leading to an amplified response and successively larger peaks of output *G*. However, this positive feedback eventually saturates, even as *I* continues to accumulate. Beyond a critical threshold, further increases in *I* have a stronger inhibitory effect on *G* than the activation mediated by *X*, leading to a progressive decline in the peak amplitude of output *G*.

Figure 4b-c shows the parameter regions for sensitization and habituation in the hybrid circuit—defined by a doubling or halving of the output peak—resemble those of the individual circuits (Figure 2c, Figure 3c). Although these regions show partial overlap, the two behaviors cannot be simultaneously maximized. This stems from their divergent parameter dependencies: while both behaviors require a low memory degradation rate *γ*, habituation strengthens monotonically with the maximal expression rate *α*, whereas sensitization weakens beyond an optimal *α* value. This inherent trade-off is a direct consequence of sharing core circuit components within a single, shared pathway.

### D. Massed–Spaced Learning

Our last example is the so-called massed–spaced learning (MSL). It refers to a cognitive strategy in which learning sessions are distributed over time (spaced learning) rather than concentrated over a short period (massed learning). This approach improves memory retention and recall by allowing time for consolidation between learning episodes. Massed–spaced learning could involve synaptic plasticity mechanisms at the single-cell level, where neurons strengthen or weaken their connections based on repeated but temporally spaced stimulation. Key processes such as long-term potentiation and synaptic tagging can be the basis for this effect, allowing individual neurons to encode and maintain learned information more effectively when spaced intervals optimize molecular and structural changes. It can be characterized on multiple scales, from synaptic changes to individual behavior, and manifests itself at both the behavioral and molecular levels (Smolen *et al*., 2016).

Although MSL is traditionally considered a neural process, a recent study suggests that similar behaviors may also be present in aneural systems, including single cells within a given tissue (Kukushkin *et al*., 2024). Their findings indicate that—under controlled conditions—these non-neural cells seem to retain information better when they are exposed to spaced intervals rather than all at once. The authors used an engineered non-neuronal reporter cell line capable of exhibiting spacing-dependent responses, offering not only increased experimental throughput to model memory formation, but also a platform to study cellular cognition outside the nervous system.

The logic architecture of MSL can be captured by a cascade of positive effects, as sketched in Figure 5a. Once again, a memory component is a requirement for learning capacity, but now the output response can be decoupled from the input events. This chain can be realized through a sequential response mechanism that involves an intermediary inducer molecule. Upon perception of a stimulus, the cell can initiate the synthesis of an intracellular inducer, which exhibits transient persistence after the stimulus ends. The MSL phenomenon can now be observed (under the right conditions) by dividing a sustained stimulus into a series of shorter stimuli separated by a delay Δ*t*, in such a way that the cell can accumulate inducer molecules over time, leading to a higher overall response. However, as will be shown below, this enhancement works only when the intervals between stimuli are shorter than the characteristic decay time of the intermediate inducer. If the delay between stimuli exceeds this critical duration, the concentration of the inducer will diminish substantially before the arrival of the subsequent stimulus, thus negating the cumulative effect.

**FIG. 5.**
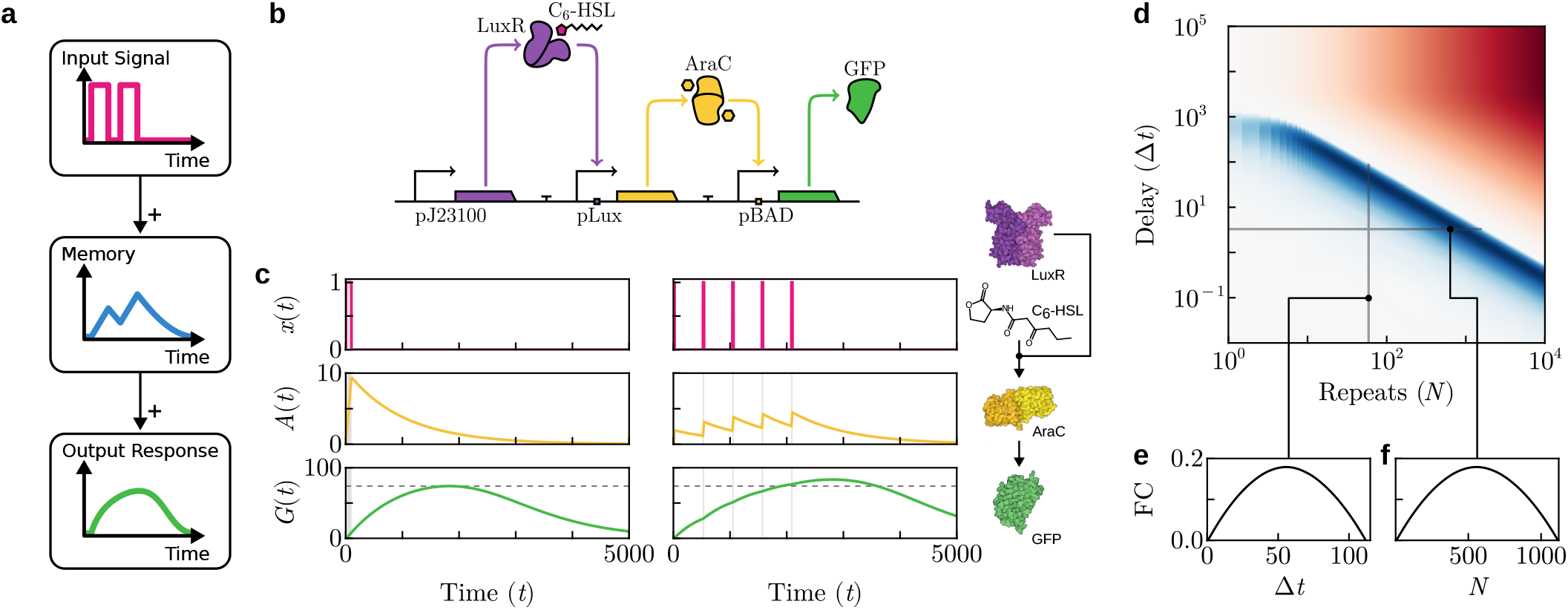
Synthetic circuit design and behavior for massed–spaced learning in a single-cell. In (a), the basic logic requirements for sensitization are outlined in a causal diagram. In (b), an explicit synthetic circuit design is shown. Receptor *X* senses the external input, which is then integrated by a downstream transcriptional activator (*A*) to drive expression of an output reporter gene (*G*). In (c), an example time-series of a response to a spaced signal with delay Δ*t* = 500 between consecutive signals, and *N* = 5 repetitions. Note that the response to the spaced signal reaches a higher maximum than that of the massed response (dashed line). In (d), the log_2_-transformed fold change (FC) of peak response with respect to the peak response for a massed signal. A signal of duration *T* = 100 is split into *N* equal pulses, separated by a delay Δ*t*. When the signal is massed (*N* = 1), the inter-pulse delay does not influence the signal, as there is no sequence of pulses to space out. The colormap is centered at FC = 0, the reference (massed peak) value, with positive deviations in blue and negative deviations in red. Slices of the Fold Change over increasing delay and increasing repetitions are displayed in the insets (e, f). Parameters: *α* = *β* = 10^−1^, *µ* = 10^−2^, *γ* = 10^−3^.

A minimal genetic implementation of this circuit, as shown in Figure 5b, consists of a linear feed-forward chain where (as in previous examples) a receptor protein *X* detects an external signal molecule *x*, leading to the production of an inducer *A*, which in turn controls the expression of the output protein *G*. Specifically, we suggest the input signal C_6_-HSL is integrated by accumulating an inducer, AraC, resulting in prolonged GFP expression that lingers after the input vanishes.

The equations governing the system are now:

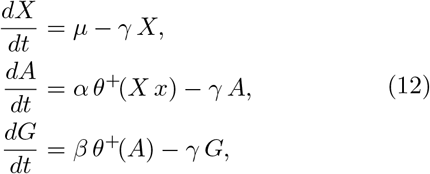

where *α, β, µ* are different promoter strengths that can be tuned separately, *γ* is the rate at which proteins degrade. The dynamical response of the circuit is summarized in the time-series in Figure 5c. For a single, continuous input pulse (left column), the intermediate activator *A* accumulates rapidly during the stimulus period, followed by an exponential decay afterwards. This transient accumulation produces a single maximum peak in the output, which quantifies the response efficiency. In contrast, when the same total input duration is divided into several evenly-spaced pulses (right column), the concentration of *A* accumulates incrementally and decays only partially during the inter-pulse intervals. This pattern of spaced reinforcement results in a higher maximum peak amplitude of the output compared to the massed case, demonstrating the expected enhancement in response. The circuit is therefore capable of distinguishing between continuous (massed) and temporally spaced input patterns, analogous to the way neural systems differentiate between massed learning and learning reinforced through repeated, spaced exposures over time.

A parameter space illustrating the relative efficiency of distributing the input into *N* evenly spaced pulses, separated by inter-pulse intervals Δ*t*, is shown in Figure 5d. The log_2_-transformed fold change in the peak response relative to the massed input case is used to quantify the enhancement due to input spacing:

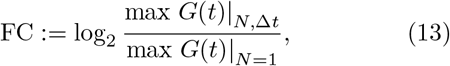

where the case *N* = 1 corresponds to massed input, and no inter-pulse delay is therefore applicable. Positive values of FC indicate an increased peak response under spaced stimulation, with the magnitude reflecting the extent of enhancement compared to the massed case. As expected, a region exists in which dividing and spacing the input pulses yields a positive gain; however, both parameters are interdependent and must be balanced.

For very short inter-pulse delays (Δ*t* → 0), the difference becomes negligible—in this limit, the signal effectively behaves as a continuous (massed) input, provided *N* remains finite and physiologically realistic.

However, if either the number of pulse repetitions or the inter-pulse delay is excessively large, the peak response decreases. This reduction arises because an excessive number of pulses (large *N*) allocates insufficient time within each pulse for the accumulation of the intermediate activator, while a prolonged inter-pulse delay (large Δ*t*) allows the activator concentration to decay significantly before the next pulse arrives.

To gain more insight into how the spacing of input pulses affects memory accumulation and downstream output, we consider a simplified version of the presented model in which each input pulse fully saturates the receptor response, and the output protein *G* is produced in an all-or-nothing manner, driven by a step activation threshold.

This can be modeled by the following simplified system:

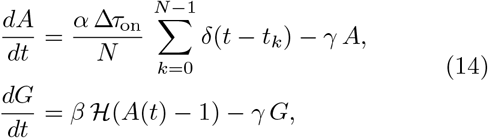

where *δ*(*t* − *t*_*k*_) denotes the Dirac delta function representing an impulsive input at discrete times *t*_*k*_ = *k* Δ*τ*_off_. Consequently, the memory variable *A* exhibits instantaneous integration of each input pulse, followed by an exponential decay in the interval before the next input arrives. Expression of output *G* is governed by a Heaviside transfer function ℋ (*A*(*t*) − 1), which acts as an abrupt switch that yields 1 when the memory is above a normal-ized threshold ℋ (*A*(*t*) ≥ 1) and 0 otherwise. Thus, production halts when the memory decays below the threshold. Let *N* identical pulses be delivered instantly at equally distant intervals of size Δ*τ* ≈ Δ*τ*_off_, with total input strength *α* Δ*τ*_on_. The state of *A* immediately after the *k*-th pulse, denoted *A*_*k*_, follows from the balance between input and decay:

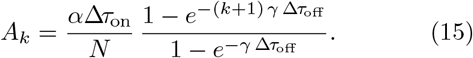

The duration of output production during the *k*-th interval, denoted 𝓁_*k*_, is determined by how long *A*(*t*) remains above the threshold, bounded by the arrival of the next pulse for all but the final interval.

Let 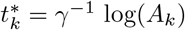 be the time expected for *A*_*k*_ *>* 1 to decay to the threshold. Then,

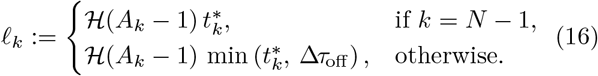

Thus, output dynamics are given by the addition of individual pulse contributions.

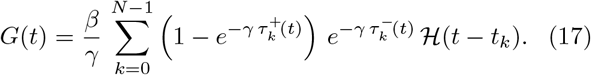

The duration of output production from the *k*-th pulse up to time *t*, denoted 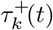,and the time elapsed since pro-duction halted, denoted 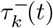,are respectively defined as:

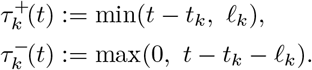

The Heaviside function ℋ (*t* − *t*_*k*_) ensures that each component only contributes after its onset time, *t*_*k*_.

With *β > γ*, the output increases during each production interval and peaks at the end of each pulse *t*_*k*_ + 𝓁_*k*_. Given equally spaced pulses and zero initial output in the first interval, each subsequent pulse starts with a positive baseline due to incomplete decay from previous inputs. This leads to a non-decreasing sequence of peak outputs. Hence, the global maximum must be at the end of the final pulse, *t* = *t*_*N*−1_ + 𝓁_*N*−1_, since no prior peak can be higher due to the cumulative effect of residual output from earlier pulses.

Under the restriction that input added in a single pulse is strong enough to sustain output production for the full relaxation phase. That is, if

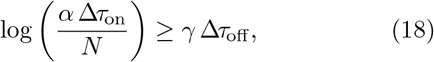

then splitting the input into *N* equally spaced pulses will yield a higher total output than a single, massed pulse. This assumption simplifies the following analysis, though it should be noted that it defines a strict subset of the parameter space where spaced pulses are advantageous. For a more detailed analysis, see Supplementary Material (SM V.A).

For a massed pulse (*N* = 1), the output production halts when 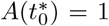, giving an output production duration:

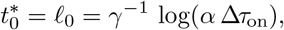

which yields a maximum output:

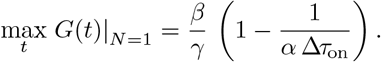

When, instead, the input is split into two equal pulses (*N* = 2), the first half-strength pulse decays for Δ*τ*_off_ until the second pulse arrives (by Assumption 18), after which the memory jumps to:

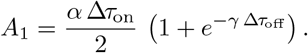

Production then halts at:

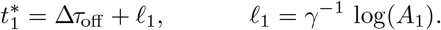

Thus, the combined active phases produce:

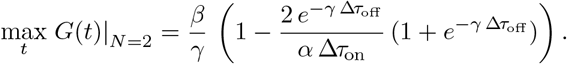

Under the previous assumptions, the region where spaced pulses are strictly better than a massed pulse is given by the inequality

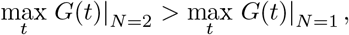

which simplifies algebraically to the condition:

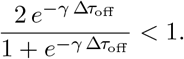

This holds when *γ* Δ*τ*_off_ > 0, ensuring that 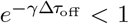. Results showing that this property also holds for the general case *N* ≥ 2 under the same assumptions are provided in Supplementary Material (SM V.A).

## III. DISCUSSION

Cognition can be defined as the acquisition, processing, storage, and use of information to generate or modulate behavior. Some single-cell organisms, from bacteria to protozoans or *Physarum* molds, are known to be able to perform a diverse range of computational tasks, including simple forms of learning. As discussed in this paper, cells within multicellular systems can also display computational tasks as part of larger integrated wholes. The study of basal cognition has deeply enlarged our comparative analysis of possible cognitions and raises some deep questions: How common is cellular cognition? How complex can it be?

A central tenet in defining cognitive complexity has to do with how organisms cope with time. When dealing with the fundamental components of basal cognition, one first aspect of this problem is how to respond to potential sources of information (from the environment or other individuals) using previous experiences. In neural systems, many of these issues are resolved at the synaptic level, whereas aneural agents have to deal with time and timing using other design principles. Circadian rhythms, for example, define a very important innovation in unicellular organisms, which internalizes night-day cycles and allows anticipation (Golden and Canales, 2003; Goldbeter, 2008; Larrondo, 2025).

In this paper, we have considered a relevant problem regarding the minimal complexity necessary for non-associative learning based on designed genetic circuits. By exploring this within the context of habituation, sensitization, and mass-paced learning, we aimed to rigorously define the key conditions that allow for their efficient implementation. A central component common to all our circuits is a memory element that provides the way to keep time and modulate the final response, directly or by means of repressor elements. Moreover, habituation and sensitization responses, as well as their combination, have been designed by means of very minor changes in transcription motifs. This suggests a high accessibility among diverse learning circuits, including potential combinations.

Our search for minimal circuits is concentrated in the use of a genetic toolkit. However, the choice of genetic motifs does not exclude other potential ways of studying and engineering cognition using other components of cellular networks. Theoretical work within systems biology, such as (Koseska and Bastiaens, 2017), has shown that biochemical networks associated with cell signaling can cause context-dependent dynamic behavior.

More generally, computational circuits can be embedded in metabolic networks (Silva-Rocha *et al*., 2011; Pandi *et al*., 2019; Carbonell *et al*., 2014).

By using synthetic biology to engineer cognitive functions, we also gain a powerful lens to investigate cellular complexity and its potential constraints. A long-term objective in this endeavor is to map the structure of the “cognitive space” in which various types of cognitive agents may reside (Solé *et al*., 2024; Solé and Seoane, 2022). This raises a fundamental question: to what extent can individual cells, whether existing independently or as part of a multicellular organism, perform cognitive tasks such as learning or anticipation? The role of gene networks in these processes may differ substantially between prokaryotic and eukaryotic systems. For example, a *Stentor* cell is approximately ∼10^5^ times larger than an *E. coli* cell, with a reproduction time of *τ*_*r*_ ≈ 2–3 days under optimal laboratory conditions, compared to the ∼2-hour replication cycle of the bacterium when grown in minimal medium. More generally, protozoan life cycles span one to three orders of magnitude longer than those of *E. coli*, allowing molecular mechanisms, including gene networks, to influence behavior throughout the life cycle of the organism. In contrast, bacterial replication times are often on the same scale as the expression dynamics of candidate synthetic circuits, constraining the integration of such mechanisms into behavior. Understanding how circuit complexity and organismal complexity are intertwined will be essential for charting the limits—and possibilities—of cellular cognition.

## ACKNOWLEDGMENTS

The authors thank the members of the Complex Systems Lab for their valuable discussions and the Santa Fe Institute, where most of this research was done. Special thanks are due to Ted Backwood for his inspiring ideas. This work has been supported by the AGAUR 2021 SGR 0075 grant and the Santa Fe Institute.

J. P. M. was supported by grant “PRE2020-091968”, funded by MCIN/AEI (10.13039/501100011033), and cofunded by the ESF through the program “Investing in your future”.

