## Supplementary Material for "Engineering Basal Cognition: Minimal Genetic Circuits for Habituation, Sensitization, and Massed–Spaced Learning"

This supplementary material extends the results presented in the main paper, where the system's learning dynamics were analyzed under a finite sequence of periodic pulses. While the finite-pulse approximation yields important insights into transient behavior and identifies relevant parameter regimes in practical scenarios, the long-term asymptotic behavior of such systems also warrants consideration.

Alternative synthetic system implementations are additionally presented, as these minimal circuits admit multiple realizations, with their own advantages and limitations.

Furthermore, some analytical approximations are derived for simplified versions of the main paper's systems. These results enable a more rigorous understanding of the underlying dynamics and support the interpretation of numerical findings.

All implementations are based on a core set of well-characterized proteins commonly used in synthetic biology. However, there are no inherent restrictions on incorporating alternative regulatory elements—such as CRISPR-based systems, conformational RNA devices like riboswitches, or gene-silencing tools such as short hairpin RNAs (shRNAs)—as long as the overall network topology remains unchanged. Integrating such elements can enhance circuit functionality, particularly by enabling regulation at the mRNA level and introducing dynamics that operate on different timescales.

For simplicity, only protein-level dynamics are taken into account, assuming that transcription and translation occur on a timescale much faster than protein degradation and functional activity. Under this assumption, mRNA levels reach a quasi-steady-state rapidly. This allows gene expression to be modeled as a single effective process, characterized by a lumped kinetic parameter that captures the combined effects of transcription and translation. This approach eliminates the need to explicitly model mRNA concentrations, significantly simplifying the system while retaining the essential features.

All circuits share the same input signal $x$, which is the N-(3-oxohexanoyl)-L-homoserine lactone (3OC6-HSL), a diffusible bacterial quorum-sensing molecule that passively crosses the cell membrane. Each circuit includes the LuxR protein ($X$), which binds the input to form an active transcription factor complex. This complex regulates promoters containing conserved DNA motifs known as *lux boxes*. The formation of the ligand-receptor complex $X$–$x$ is modeled as a first-order interaction, proportional to the product of the concentrations: $X–x \propto X\,x$. In all cases except one, the complex acts as an activator of transcription.

Additional transcription factors used include two repressor proteins—LacI ($I$) and TetR ($R$)—and one activator, AraC ($A$).

The circuit's output $G$ will be a green fluorescent protein (GFP), which would enable straightforward experimental validation and real-time monitoring of dynamic responses *in vivo*. However, additional functionality can be incorporated by co-expressing the fluorescent protein within the same operon as the target effector protein, enabling simultaneous expression and tracking.

---

Transcriptional regulation by these factors is modeled using Hill-type transfer functions. For a repressor and an activator, respectively:

$$\theta_K^-(I) := \frac{1}{1+\left(\frac{I}{K}\right)^2}, \qquad \theta_K^+(X\,x) := \frac{\left(\frac{X\,x}{K}\right)^2}{1+\left(\frac{X\,x}{K}\right)^2}.$$

In most cases, a normalized version with the half-saturation constant $K_{1/2} = 1$ is used, simplifying the expressions by omitting the subscript $K$:

$$\theta^-(I) := \frac{1}{1+I^2}, \qquad \theta^+(X\,x) := \frac{(X\,x)^2}{1+(X\,x)^2}.$$

To establish a separation of timescales—critical for dynamic control and memory formation—fast-degrading variants of proteins can be generated by appending short peptide sequences that target the protein for degradation by the cell's proteolytic machinery. This degradation tag effectively reduces the protein's half-life. For simplicity, a suffix "-LVA" is used to denote proteins containing such a tag, where the LVA sequence is a widely used degradation tag in synthetic biology. It is derived from the ssrA tag and directs the tagged protein for recognition and rapid degradation by protease complexes such as ClpXP in *E. coli* and other bacterial hosts.

As a measure of habituation and sensitization strength, the fold change, FC, defined as the $\log_2$ ratio between the maximum response after the final stimulus and the peak response observed throughout the simulation time span will be used:

$$\mathrm{FC} := \log_2 \frac{\max G(\tau)}{\max G(T)}, \qquad \tau \in [t_{n-1}, t_n). \tag{1}$$

### I. HABITUATION

#### A. Habituation with an Incoherent Feed-Forward Loop

Taking the system for habituation presented in the main paper (see Figure 1a), in which a constitutively expressed receptor $X$, upon binding the input signal $x$, acts as a transcriptional activator for the repressor $I$. Concurrently, the output ($G$) is expressed from a promoter that is dual-regulated: it is activated by the $X$–$x$ complex and repressed by $I$. As a result, $G$ is produced only when $x$ is present, but its maximum expression rate decreases as $I$ accumulates.

To establish a separation of timescales—ensuring that only the repressor $I$ accumulates over successive input pulses—the output protein $G$ is fused to the LVA degradation tag (GFP-LVA), resulting in a short half-life and preventing its buildup between pulses. For compatibility with other circuits discussed later, the receptor $X$ is also assumed to carry the LVA tag (LuxR-LVA). However, since $X$ remains at steady state throughout the dynamics due to constitutive expression, its functional behavior can be equivalently reproduced by removing the degradation tag and adjusting the promoter strength to maintain the same effective concentration.

The following equations are used for modeling the system:

$$\begin{aligned} \frac{dX}{dt} &= \mu - \lambda\,X, \\ \frac{dI}{dt} &= \alpha\,\theta^+(X\,x) - \gamma\,I, \\ \frac{dG}{dt} &= \beta\,\theta^+(X\,x)\,\theta^-(I) - \lambda\,G, \end{aligned} \tag{2}$$

which is a realization of an Incoherent Feed-Forward loop (I-FFL). Key parameters are $\alpha$, $\beta$, and $\mu$, represent the maximum promoter expression rates which can be independently tuned. Basal degradation rate $\gamma$ for untagged proteins and increased rate $\lambda$ for tagged proteins, with $\lambda \geq \gamma$.

Habituation is most effective when the memory degradation rate ($\gamma$) is low. As it can be seen in Figure 1b, if memory degradation is high, habituation won't occur because the accumulated repressor $I$ vanishes before the next stimulus

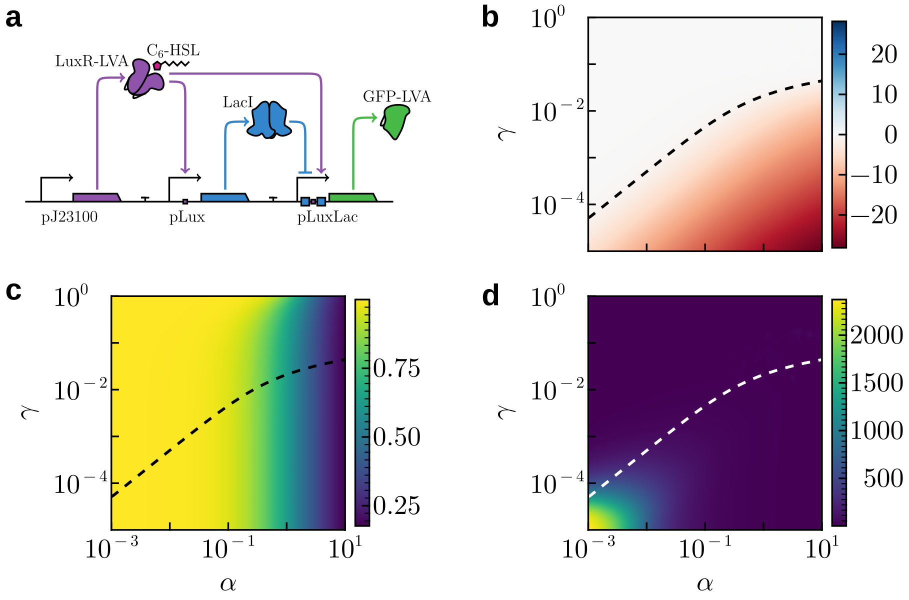


FIG. 1 **Habituation with an incoherent feed-forward loop.** (a) The proposed genetic circuit implementing the habituation incoherent feed-forward loop motif. (b) Long-term parameter space where habituation occurs, showing the $\log_2$-transformed fold change between the long-term peak magnitude and the first peak. The dashed line at $-1$ indicates a halving in the long-term peak magnitude compared to the first peak. (c) Magnitude of the first response peak, representing the initial maximal output of the system before habituation effects become significant. (d) Number of peaks required for the system's response to enter an $\epsilon$-ball ($\epsilon = 10^{-3}$) around the expected steady-state peak magnitude, where subsequent fluctuations remain within this neighborhood. Parameters: $\beta = 2$, $\mu = 1$, $\lambda = 1$.

arrives. When degradation is slow, increasing the maximum promoter expression for the repressor ($\alpha$) leads to a greater magnitude of fold change (FC) between the first and last peaks. This means the system habituates more strongly. At the same time, when the repressor builds up faster ($\alpha \gg \gamma$), the initial response (without any previous input) also decreases, as shown in Figure 1c.

There are inherent limits to how much the output can be decreased. These can be approximated by finding the steady-state values for a sustained input, $x$.

The steady-state equations for the system are as follows:

$$\begin{aligned} \frac{dX}{dt} &= 0 \implies X^* = \frac{\mu}{\lambda}, \\ \frac{dI}{dt} &= 0 \implies I^* = \frac{\alpha}{\gamma} \frac{(X^* x)^2}{1 + (X^* x)^2}, \\ \frac{dG}{dt} &= 0 \implies G^* = \frac{\beta}{\lambda} \frac{(X^* x)^2}{1 + (X^* x)^2} \frac{1}{1 + (I^*)^2}. \end{aligned}$$

Substituting $X^* = \frac{\mu}{\lambda}$, the steady-state values become:

$$I^* = \frac{\alpha}{\gamma} \frac{\left(\frac{\mu x}{\lambda}\right)^2}{1 + \left(\frac{\mu x}{\lambda}\right)^2}, \qquad G^* = \frac{\beta}{\lambda} \frac{\left(\frac{\mu x}{\lambda}\right)^2}{1 + \left(\frac{\mu x}{\lambda}\right)^2} \frac{1}{1 + (I^*)^2},$$

which when fully habituated ($x \to \infty$) becomes:

$$I^* \approx \frac{\alpha}{\gamma}, \qquad G^* \approx \frac{\beta}{\lambda} \frac{1}{1 + \left(\frac{\alpha}{\gamma}\right)^2} = \frac{\beta \gamma^2}{\lambda(\alpha^2 + \gamma^2)}. \tag{3}$$

This last expression for $G^*$ provides an upper bound on the maximum repression of the output.

Following Equations (3), it is possible to maintain the strength of habituation by scaling down both production and degradation accordingly. This avoids the problem mentioned earlier where the first response is already habituated (see Figure 1c). This approach, however, presents a trade-off. Slowing down the memory dynamics means a higher number of peaks are required for the system to habituate. This can become realistically inefficient as the goal is to respond only after a small number of peaks have been presented (see Figure 1d).

#### B. Habituation with a Negative Self-Feedback Loop

An alternative design for a habituating circuit uses a single component: a receptor that can act as both an activator and a repressor. While wild-type LuxR primarily functions as an activator, but certain promoters can enable LuxR-mediated repression, making this design a possibility.

In this system, receptor $X$ (LuxR) activates the output while also repressing its own expression. The circuit's output $G$ (GFP-LVA)in principle, should be produced only when the input signal $x$ is present, and its expression rate should reduced over time by the accumulation of its own repressor form.

However, a significant limitation of this design arises when assuming the Hill-like dynamics for transcriptional regulatiors: the repressor concentration required for effective self-inhibition far exceeds the activator concentration needed to substantially reduce output expression. This imbalance makes it challenging to tune the system for effective self-inhibition, as the repressor needs to accumulate to a very high level to significantly affect its own production while at this steady state concentration, output still is produced.

The following equations are used for modeling the system:

$$\begin{aligned} \frac{dX}{dt} &= \alpha\, \theta^{-}_{K^{-}}(X\, x) - \gamma\, X, \\ \frac{dG}{dt} &= \beta\, \theta^{+}_{K^{+}}(X\, x) - \lambda\, G, \end{aligned} \tag{4}$$

where the interesting parameters in this case are the the activation and repression thresholds ($K^{+}$ and $K^{-}$).

This negative self-feedback architecture differs from the incoherent feed-forward loop in that it relies on a single regulatory component ($X$) to both drive and suppress output expression. In practice, this means that for the system to exhibit robust habituation, the repressive effect of $X$ must dominate at high concentrations, while still allowing sufficient initial activation when the input signal $x$ is first detected. However, as transcriptional repression often requires much higher protein concentrations than activation, the dynamic range of the system may be insufficient to achieve meaningful output suppression without severely compromising initial responsiveness. Additionally, unlike the I-FFL, where a repressor accumulates independently of the receptor $X$, this self-regulating motif ties repression directly to the same molecule responsible for activation. As a result, tuning the system for effective habituation becomes more challenging, as adjustments to promoter strengths or degradation rates affect both activation and repression simultaneously. This interdependence limits the flexibility of parameter optimization compared to the more modular design of the feed-forward loop.

Based on the model's behavior shown in Figure 2b, a noticeable decrease in the fold change between the first and final peak only occurs when the activation threshold ($K^{+}$) is orders of magnitude higher than the repression threshold ($K^{-}$). Even then, the reduction is modest, never reaching a halving of the initial response. This need level of parameter tuning also severely compromises the system's initial responsiveness, as the maximum output is significantly lowered (Figure 2c). As a result, this regulatory motif is largely impractical; it requires an unrealistic difference in its half-saturation constants, yields a low maximum output, and offers only minimal habituation.

#### C. Habituation with a Negative Feedback Loop

The issues seen in the negative self-feedback loop can be partially addressed by introducing an intermediate repressor as memory component, with no change in the circuit's topology. This design, shown in Figure 3a, separates the roles of activation and repression into distinct molecular components. This independence enables a more robust and flexible system where the intermediate repressor can build up over successive inputs, effectively creating a cellular memory of previous stimuli.

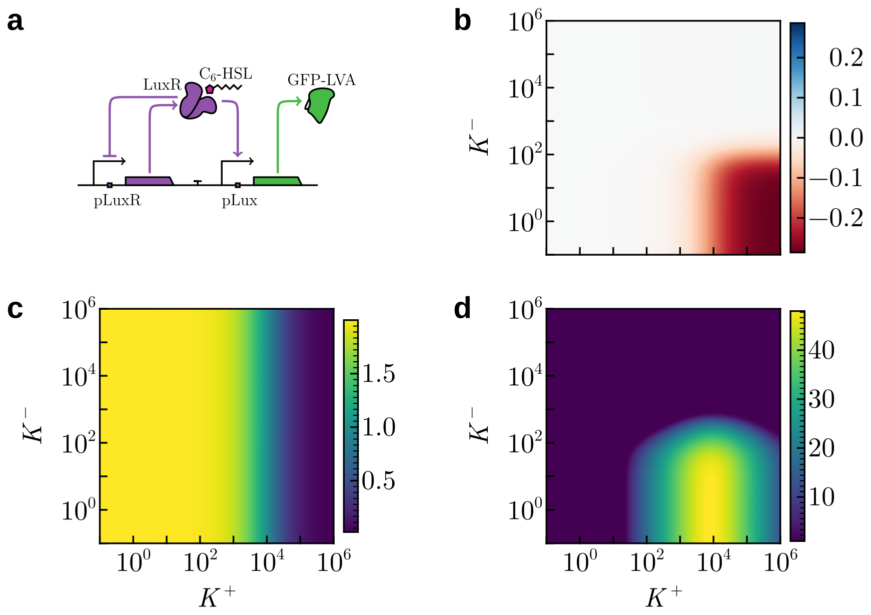


FIG. 2 **Habituation with a negative self-feedback loop.** (a) The proposed genetic circuit implementing the habituation with negative autoregulation. (b) Long-term parameter space where habituation occurs, showing the $\log_2$-transformed fold change between the long-term peak magnitude and the first peak. No dashed line is present as there is not enough habituation to halve the first peak. (c) Magnitude of the first response peak, representing the initial maximal output of the system before habituation effects become significant. (d) Number of peaks required for the system's response to enter an $\epsilon$-ball ($\epsilon = 10^{-3}$) around the expected steady-state peak magnitude, where subsequent fluctuations remain within this neighborhood. Parameters: $\alpha = 10^{-1}$, $\beta = 2$, $\gamma = 10^{-3}$, $\lambda = 1$.

The following equations are used for modeling the system:

$$\begin{aligned} \frac{dX}{dt} &= \mu\,\theta^{-}(I) - \lambda\,X, \\ \frac{dI}{dt} &= \alpha\,\theta^{+}(X\,x) - \gamma\,I, \\ \frac{dG}{dt} &= \beta\,\theta^{+}(X\,x) - \lambda\,G, \end{aligned} \tag{5}$$

with key parameters $\alpha$, $\beta$, and $\mu$, represent the maximum promoter expression rates which can be independently tuned. Basal degradation rate $\gamma$ for untagged proteins and increased rate $\lambda$ for tagged proteins, with $\lambda \geq \gamma$.

Given a sustained input $x$, the steady state can be derived as follows:

$$\begin{aligned} \frac{dX}{dt} = 0 &\implies X^* = \frac{\mu}{\lambda}\frac{1}{1+(I^*)^2}, \\ \frac{dI}{dt} = 0 &\implies I^* = \frac{\alpha}{\gamma}\frac{(X^*\,x)^2}{1+(X^*\,x)^2}, \\ \frac{dG}{dt} = 0 &\implies G^* = \frac{\beta}{\lambda}\frac{(X^*\,x)^2}{1+(X^*\,x)^2}. \end{aligned}$$

For the case of strong input ($x \to \infty$), the steady-state values approximate:

$$X^* \approx \frac{\mu}{\lambda}\frac{1}{1+\left(\frac{\alpha}{\gamma}\right)^2}, \qquad I^* \approx \frac{\alpha}{\gamma}, \qquad G^* \approx \frac{\beta}{\lambda}. \tag{6}$$

Comparing the steady-state output $G^*$ of the fully habituated system to the I-FFL case from steady-state solutions given in Equation (3), note that the I-FFL consistently achieves a greater long-term repression, by a factor:

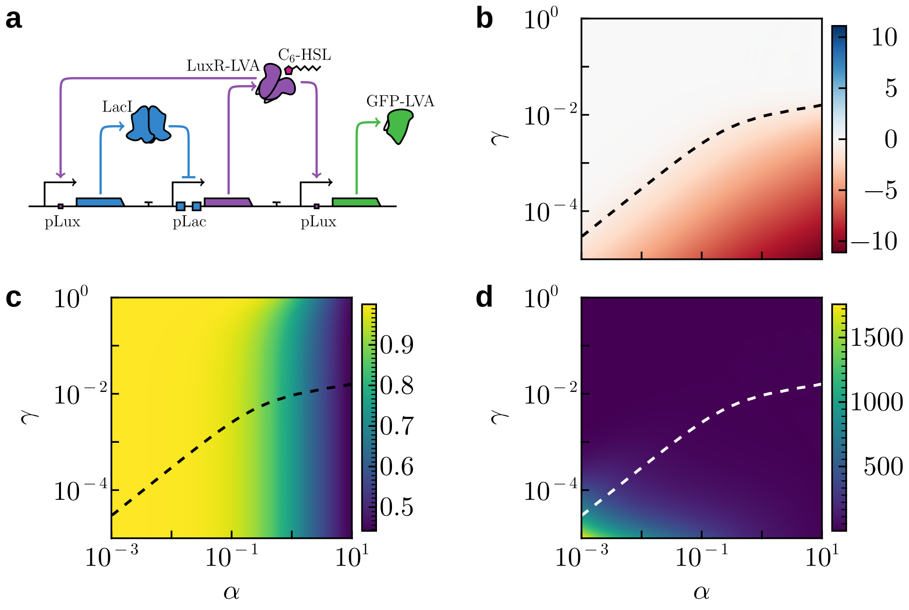


FIG. 3 **Habituation with a negative feedback.** (a) The proposed genetic circuit implementing habituation with a negative feedback through accumulation of an intermediate repressor. (b) Long-term parameter space where habituation occurs, showing the $\log_2$-transformed fold change between the long-term peak magnitude and the first peak. The dashed line at $-1$ indicates a halving in the long-term peak magnitude compared to the first peak. (c) Magnitude of the first response peak, representing the initial maximal output of the system before habituation effects become significant. (d) Number of peaks required for the system's response to enter an $\epsilon$-ball ($\epsilon = 10^{-3}$) around the expected steady-state peak magnitude, where subsequent fluctuations remain within this neighborhood. Parameters: $\beta = 2$, $\mu = 1$, $\lambda = 1$.

The I-FFL case consistently achieves greater long-term repression than the fully habituated system using a negative feedback. This can be demonstrated by comparing the solutions from Equations (3) to Equations (6), which shows the I-FFL motif can decrease the output concentration by a larger multiplicative factor

$$\frac{\gamma^2}{(\alpha^2 + \gamma^2)} < 1.$$

Similar to the I-FFL case, the memory degradation rate ($\gamma$) must remain low to ensure that memory persists across stimulus pulses (Figure 3b). When degradation is slow, increasing the maximum promoter expression of the repressor ($\alpha$) results in a larger fold change (FC) between the first and last response peaks; however, this comes at the cost of a reduced initial response, as illustrated in Figure 3c.

The same trade-off for arbitrarily small values of $\alpha$ and $\gamma$ also holds, where full habituation demands a number of stimulus peaks that becomes physiologically unrealistic (Figure 3d).

### II. MEMORY ACCUMULATION

Given the central role played by the memory component in shaping the dynamics of the learning motifs explored in this work, it becomes essential to characterize the conditions under which such a memory can accumulate over time. This requires a detailed analysis of the parameter space governing the interplay between memory induction and decay, particularly in response to periodic external stimuli. To this end, we present a formal and extended treatment of the system's dynamics, offering a comprehensive approximation of the regime in which sustained memory accumulation is feasible.

Consider a scenario in which the temporal evolution of the memory component—denoted as $I(t)$—is driven by a sequence of repeated external inputs. These inputs are structured in a periodic fashion, with each cycle consisting of two distinct phases: an active phase of duration $\Delta\tau_{\text{on}}$, during which the memory-inducing stimulus is applied, followed by a relaxation phase of duration $\Delta\tau_{\text{off}}$, during which no external input is present. The total period of this repeating signal is therefore given by $\Delta\tau = \Delta\tau_{\text{on}} + \Delta\tau_{\text{off}}$. This temporal structure mimics patterns of neural or cognitive activation, where brief periods of stimulation are interspersed with longer recovery intervals.

During the active phase of duration $\Delta\tau_{\text{on}}$, the memory component $I(t)$ is subject to a constant production process, increasing at a rate $\alpha$. Concurrently, across both the active and relaxation phases, the component undergoes exponential decay at a constant rate $\gamma$, reflecting the natural forgetting or dissipation of memory in the absence of reinforcement.

Under the assumption that the duration of the active phase is much shorter than that of the subsequent relaxation phase (i.e., $\Delta\tau_{\text{on}} \ll \Delta\tau_{\text{off}}$) the input can be effectively approximated as an instantaneous pulse. This simplification approximates the repeated stimulation as discrete, impulsive events separated by extended intervals of passive decay. Consequently, the overall dynamics of the memory component can be described by a piece-wise differential equation:

$$\frac{dI}{dt} = \alpha\, \Delta\tau_{\text{on}} \sum_{i=1}^{N} \delta(t - t_i) - \gamma\, I, \tag{7}$$

where $\delta(t - t_k)$ denotes the Dirac delta function representing an impulsive input at discrete times $t_k = k\ \Delta\tau$, with $k = 1, \ldots, N$ indexing the stimulus cycles. The prefactor $\alpha\, \Delta\tau_{\text{on}}$ captures the total memory increment delivered during each brief active phase, effectively modeling the integrated effect of a small, finite stimulus as an instantaneous "jump" of magnitude $\alpha\, \Delta\tau_{\text{on}}$.

The conditions under which the memory component $I(t)$ can grow over time can be analyzed analytically under the assumption that memory does not saturate its downstream effects—specifically, $I(t) \ll \alpha/\gamma$—throughout the system's evolution. This linear regime assumption allows to neglect nonlinear saturation effects and focus on the cumulative dynamics driven by repeated stimulation.

Begin with the general solution to the differential equation (7), which governs the evolution of $I(t)$ under impulsive inputs:

$$I(t) = I_0\, e^{-\gamma\, t} + \alpha\, \Delta\tau_{\text{on}} \sum_{i=1}^{N} \mathcal{H}(t - t_i)\, e^{-\gamma\,(t - t_i)},$$

where $t_k = k\, \Delta\tau$ denotes the time of the $k$-th stimulus, and $\mathcal{H}(t - t_k)$ is the Heaviside step function, indicating that each impulse contributes only for $t \geq t_k$. The first term represents the decay of the initial memory $I_0$, which becomes negligible over long times.

The Heaviside function

$$\mathcal{H}(u) = \begin{cases} 0 & \text{for } u < 0, \\ 1 & \text{for } u \geq 0, \end{cases} \tag{8}$$

is the antiderivative of the Dirac delta $\delta(u)$:

$$\frac{d}{du}\mathcal{H}(u) = \delta(u) \iff \mathcal{H}(u) = \int_{-\infty}^{u} \delta(s)\, ds.$$

This expresses that the delta function models an instantaneous impulse at $u = 0$, while the Heaviside function represents its cumulative effect, switching from 0 to 1 at that point.

When the initial conditions are zero, or for long-term dynamics ($t \to \infty$), the initial memory component decays exponentially and becomes negligible, leaving only the sustained contribution from repeated inputs:

$$I(t) = \alpha\, \Delta\tau_{\text{on}} \sum_{i=1}^{\infty} \mathcal{H}(t - t_i)\, e^{-\gamma\,(t - t_i)}. \tag{9}$$

Consider a time $t$ located in the $n$-th cycle: $t = n\, \Delta\tau + \tau$, where $0 \leq \tau < \Delta\tau$. At this point, all stimuli up to and including the $n$-th have occurred. Substituting into the expression above:

$$I(t) = \alpha\, \Delta\tau_{\text{on}} \sum_{i=1}^{n} e^{-\gamma\,(n\,\Delta\tau + \tau - i\,\Delta\tau)} = \alpha\, \Delta\tau_{\text{on}}\, e^{-\gamma\,\tau} \sum_{i=1}^{n} e^{-\gamma\,(n-i)\,\Delta\tau}.$$

By changing the summation index to $k = n - i$, the expression becomes:

$$I(t) = \alpha\, \Delta\tau_{\text{on}}\, e^{-\gamma\, \tau} \sum_{k=0}^{n-1} \left(e^{-\gamma\, \Delta\tau}\right)^k ,$$

which is a finite geometric series with ratio $r = e^{-\gamma\, \Delta\tau}$. Its sum is given by:

$$\sum_{k=0}^{n-1} r^k = \frac{1 - r^n}{1 - r} = \frac{1 - e^{-\gamma\, n\, \Delta\tau}}{1 - e^{-\gamma\, \Delta\tau}}.$$

Thus, the memory accumulated up to time $t = n\, \Delta\tau + \tau$ can be expressed as:

$$I(t) = \alpha\, \Delta\tau_{\text{on}}\, e^{-\gamma\, \tau}\, \frac{1 - e^{-\gamma\, n\, \Delta\tau}}{1 - e^{-\gamma\, \Delta\tau}}. \tag{10}$$

In the long-time limit, $n \to \infty \implies e^{-\gamma\, n\, \Delta\tau} \to 0$, and the memory settles into a periodic steady state:

$$I(t) \approx \alpha\, \Delta\tau_{\text{on}}\, \frac{e^{-\gamma\, \tau}}{1 - e^{-\gamma\, \Delta\tau}}, \qquad \tau \in [0, \Delta\tau). \tag{11}$$

This expression describes a memory trace that oscillates periodically within each cycle, decaying exponentially between inputs and jumping instantaneously at each stimulus onset. The amplitude of oscillation is bounded by:

- Immediately after a stimulus ($\tau \to 0^+$):

$$I_{\text{max}} := \lim_{\tau \to 0^+} I(t) \approx \frac{\alpha\, \Delta\tau_{\text{on}}}{1 - e^{-\gamma\, \Delta\tau}},$$

- Just before the next stimulus ($\tau \to \Delta\tau^-$):

$$I_{\text{min}} := \lim_{\tau \to \Delta\tau^-} I(t) \approx \frac{\alpha \Delta\tau_{\text{on}}\, e^{-\gamma \Delta\tau}}{1 - e^{-\gamma \Delta\tau}}.$$

The discontinuity at each impulse corresponds to the instantaneous addition of memory $\alpha\, \Delta\tau_{\text{on}}$. This is consistent with the model, as:

$$I(n\, \Delta\tau^+) - I(n\, \Delta\tau^-) = \alpha\, \Delta\tau_{\text{on}} \left(\frac{1 - e^{-\gamma\, \Delta\tau}}{1 - e^{-\gamma\, \Delta\tau}}\right) = \alpha\, \Delta\tau_{\text{on}},$$

confirming that the jump magnitude matches the input size.

Furthermore, in the limit of slow decay relative to the period ($\gamma\, \Delta\tau \ll 1$), the following expansion is valid:

$$1 - e^{-\gamma\, \Delta\tau} \approx \gamma\, \Delta\tau,$$

leading to an even further simplified expression for long times:

$$I(t) \approx \frac{\alpha\, \Delta\tau_{\text{on}}}{\gamma\, \Delta\tau_{\text{off}}}\, e^{-\gamma\, \tau}, \qquad \tau \in [0, \Delta\tau).$$

### III. SENSITIZATION

#### A. Sensitization with a Positive Feedback Loop

Taking the system for sensitization presented in the main paper (see Figure 4a), the architecture is modified from the habituation circuit from Figure 1a. In this design, receptor $X$ (LuxR-LVA) is no longer under constitutive control but is instead repressed by a new regulator, $R$ (TetR-LVA). This introduces a key feedback mechanism: the repressor $I$ (LacI), which is transcriptionally activated by the $X$–$x$ complex as before, now acts as a repressor for $R$. Consequently,

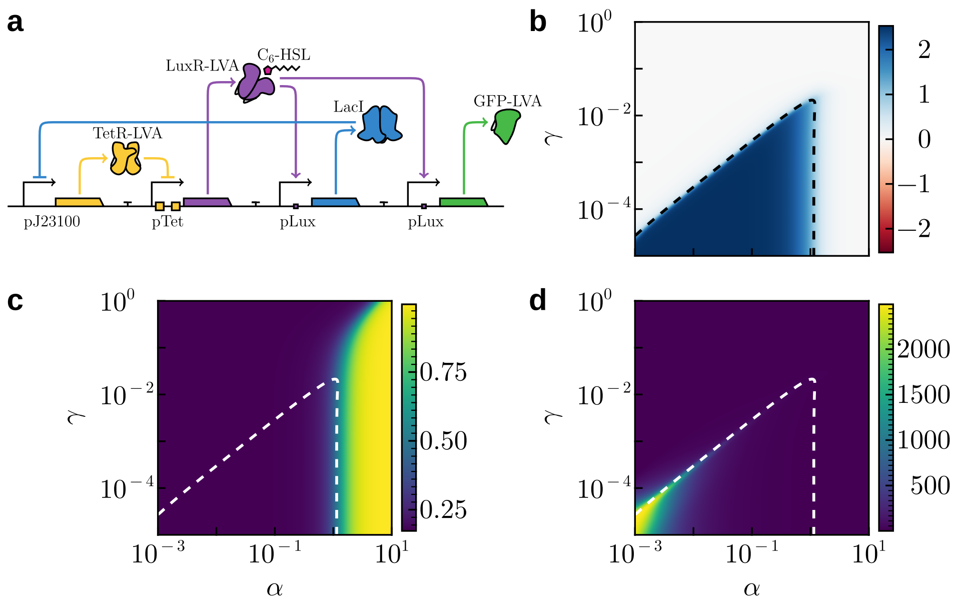


FIG. 4 **Sensitization with a Positive Feedback Loop.** (a) The proposed genetic circuit implementing the sensitization positive feedback loop motif using a chain of repressors. (b) Long-term parameter space where sensitization occurs, showing the $\log_2$-transformed fold change between the long-term peak magnitude and the first peak. The dashed line at 1 indicates a doubling in the long-term peak magnitude compared to the first peak. (c) Magnitude of the first response peak, representing the initial maximal output of the system before sensitization effects become significant. (d) Number of peaks required for the system's response to enter an $\epsilon$-ball ($\epsilon = 10^{-3}$) around the expected steady-state peak magnitude, where subsequent fluctuations remain within this neighborhood. Parameters: $\beta = 2$, $\mu = 1$, $\lambda = 1$, $\rho = 1.5$.

the expression of the receptor $X$ is regulated through a chain of two repressors, creating a form of transcriptional positive feedback.

The output $G$ (GFP-LVA) is expressed from a promoter that is activated directly by the $X$–$x$ complex, without repression from $I$. Thus, production of $G$ is dependent on the presence of the input signal and the available receptor concentration.

The key to the memory effect is that only the repressor $I$ has a slow degradation rate ($\gamma$). This allows $I$ to accumulate and persist long after the input signal is gone. Its continued presence represses $R$, which in turn disinhibits the production of new $X$ receptors. The fast degradation ($\lambda$) of both $X$ and $R$ ensures their concentrations rapidly reach a quasi-steady-state set by the slowly changing concentration of $I$.

This system is modeled by the following equations:

$$\begin{aligned}
\frac{dX}{dt} &= \mu\,\theta^{-}(R) - \lambda\,X,\\
\frac{dI}{dt} &= \alpha\,\theta^{+}(X\,x) - \gamma\,I,\\
\frac{dR}{dt} &= \mu\,\theta^{-}(I) - \lambda\,R,\\
\frac{dG}{dt} &= \beta\,\theta^{+}(X\,x) - \lambda\,G,
\end{aligned} \tag{12}$$

where $\mu$, $\alpha$, and $\beta$ are maximum promoter expression, which can be independently tuned to shape the system's dynamics. Basal degradation rate $\gamma$ for untagged proteins and increased rate $\lambda$ for tagged proteins, with $\lambda \geq \gamma$.

Sensitization requires slow memory degradation ($\gamma$). If degradation is too fast, the repressor protein $I$ degrades before the next stimulus arrives, preventing sensitization (Figure 4b).

When degradation is slow, a higher repressor production rate $\alpha$) increases the fold change FC) up to an optimal point. If $\alpha$ is too low, insufficient repressor $I$ accumulates and thus no receptor $X$ can be expressed to effectively activate the output promoter. Conversely, if $\alpha$ is too high, the system's response to a short input pulse is already near its maximum possible response. Since the system's output is bounded, this high initial response limits the potential fold change for subsequent stimuli. This explains the sharp drop in FC observed at high $\alpha$ values as the first observed peak rapidly increases (see Figure 4c).

Beyond these constraints, the parameter space for robust sensitization is further limited by the requirement for a realistic number of priming pulses. For certain parameter combinations, while sensitization is a possibility, it demands an impractical number of stimuli (see Figure 4d).

#### B. Sensitization with Self-activation Loop

While the sensitization systems presented before are quite minimal, there is an even more simple motif which can lead to a sensitized response over time: a self-activation loop in the receptor.

This synthetic circuit design is based on an activator mediated by a small molecule $X$, in this example, the Lux receptor LuxR, which upon binding an external input signal $x$—the QS molecule$C_6$-HSL—binds to its own promoter to activate it and accumulate more receptor, which would allow the cell to respond more to subsequent input presentations. The output signal $G$ is the fluorescent protein GFP-LVA as used in the previous examples. Both $X$ and $G$ are expressed under an inducible promoter regulated by the $X$–$x$ complex.

The equations governing the system are as follows:

$$\frac{dX}{dt} = \alpha\, \left[\varepsilon + (1-\varepsilon)\, \theta^{+}(X\, x)\right] - \gamma\, X, \tag{13}$$

$$\frac{dG}{dt} = \beta\, \left[\varepsilon + (1-\varepsilon)\, \theta^{+}(X\, x)\right] - \lambda\, G, \tag{14}$$

where $\alpha$, and $\beta$ are maximum promoter expression, which can be independently tuned to shape the system's dynamics. Basal degradation rate $\gamma$ for untagged protein $X$ and increased rate $\lambda$ for tagged output protein $G$, with $\lambda \geq \gamma$.

The basal expression $\varepsilon$ is necessary to avoid an inactive fixed point at $X^* = 0$, from which the system cannot escape as it would not able to sense the external input. An alternative way to solve this problem would be to add an additional source for $X$ under a weak constitutive promoter (leading to a rescaled system), but in any case this is not the only problem this two-component system presents.

While theoretically the system works and there is a region in the parameter space where sensitization can be observed (Figure 5b), the viable parameter range for this behavior is far narrower than in the alternative circuits. This range is also highly sensitive to the basal expression rate $\varepsilon$: it must be high enough to kickstart the positive feedback mechanism but low enough that the initial response is not saturated, leaving room for a stronger, sensitized response later.

#### C. Sensitization with a Coherent Feed-Forward Loop

Another possible implementation for a simple sensitization circuit is presented in Figure 6a. In this design, the receptor $X$ (LuxR-LVA) is constitutively expressed, providing a constant baseline level of signal detection. The input signal $x$, upon binding to $X$, activates the expression of repressor $R$ (TetR). Protein $R$ then acts to repress a second repressor, $I$ (LacI-LVA).

The output of the system, $G$ (GFP-LVA), is expressed under a hybrid promoter designed to be active only under two conditions: the presence of the activating $X$–$x$ complex and the absence of the repressor $I$. This topology creates a coherent type-1 feed-forward loop (C1-FFL), in which the input signal regulates the output through two parallel, positive pathways. The first is a direct path via transcriptional activation by the $X$–$x$ complex. The second is an indirect path where the input activates $R$, which in turn represses $I$. This double-negative logic (repression of a repressor) results in a net positive activation of the output $G$.

The key in the memory mechanism relies on a separation of timescales, created by the distinct degradation rates of the system's components. Repressor $R$ degrades slowly, allowing it to accumulate over successive input pulses. Its persistent presence ensures the continuous repression of the fast-degrading intermediate repressor $I$. As a result, the concentration of $I$ rapidly depletes and reaches a quasi-steady-state set by the slowly changing concentration

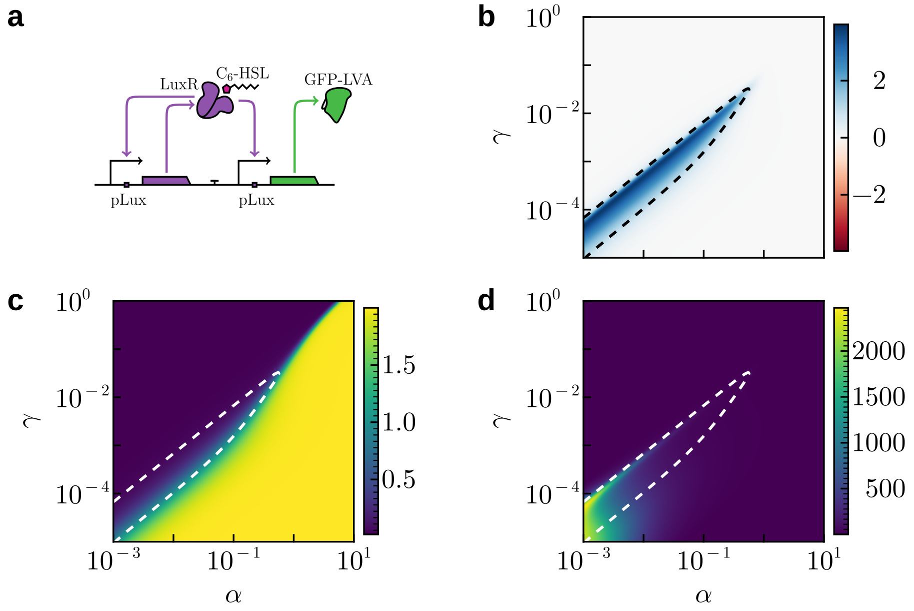


FIG. 5 **Sensitization with Self-activation Loop.** (a) The proposed genetic circuit implementing the sensitization positive feedback loop motif using a self-activation loop. (b) Long-term parameter space where sensitization occurs, showing the $\log_2$-transformed fold change between the long-term peak magnitude and the first peak. The dashed line at 1 indicates a doubling in the long-term peak magnitude compared to the first peak. (c) Magnitude of the first response peak, representing the initial maximal output of the system before sensitization effects become significant. (d) Number of peaks required for the system's response to enter an $\epsilon$-ball ($\epsilon = 10^{-3}$) around the expected steady-state peak magnitude, where subsequent fluctuations remain within this neighborhood. Parameters: $\beta = 2$, $\lambda = 1$, $\varepsilon = 10^{-2}$.

of $R$. This disinhibition of the output promoter, caused by the slow depletion of $I$, is what leads to the sensitized response.

This system is modeled by the following equations:

$$\begin{aligned}\frac{dX}{dt} &= \mu - \lambda X,\\ \frac{dR}{dt} &= \alpha\,\theta^{+}(X\,x) - \gamma\,R,\\ \frac{dI}{dt} &= \rho\,\theta^{-}(R) - \lambda\,I,\\ \frac{dG}{dt} &= \alpha\,\beta\,\theta^{+}(X\,x)\,\theta^{-}(I) - \lambda\,G,\end{aligned} \tag{15}$$

where $\alpha$, $\beta$, $\mu$, $\rho$ are maximum promoter expression rates, which can be independently tuned to shape the system's dynamics. The separation of timescales is enforced by the basal degradation rate $\gamma$ for the slow memory protein $R$ and the increased rate $\lambda$ for the all other proteins, with $\lambda \geq \gamma$.

The requirements for achieving sensitization in CFF motifs are fundamentally similar to those in the positive feedback case, as both rely on accumulation of a slow-degrading memory component. Thus, sensitization requires slow memory degradation ($\gamma$). If degradation is too fast, the repressor protein $R$ degrades before the next stimulus arrives, preventing sensitization (Figure 6b).

When the degradation rate is slow, the fold change (FC) increases with the repressor production rate ($\alpha$) up to an optimal value. If $\alpha$ is too low, insufficient repressor ($R$) accumulates, leaving the inhibitor ($I$) active and preventing input-mediated activation of the output promoter. Conversely, if $\alpha$ is too high, the system's strong initial response to a short input pulse depletes $I$ below its functional concentration, causing the output to approach its maximum. Because the output is bounded, this prematurely saturated response diminishes the potential fold change for subsequent stimuli. This accounts for the sharp decline in FC at high $\alpha$, where the first response peak is rapidly maximized (see Figure 6c).

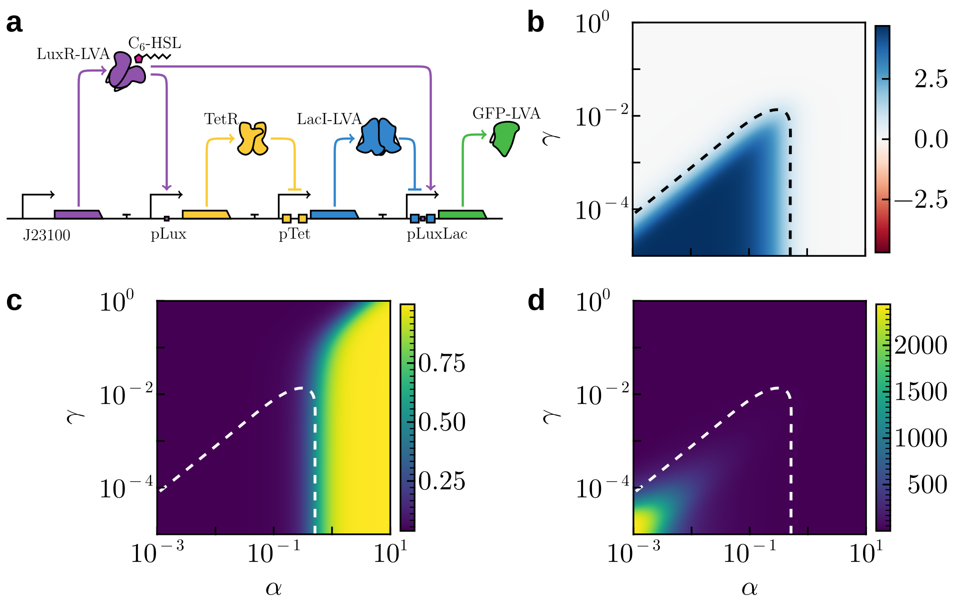


FIG. 6 **Sensitization with a Coherent Feed-Forward Loop.** (a) The proposed genetic circuit implementing the sensitization coherent feed-forward loop motif using a chain of repressors. (b) Long-term parameter space where sensitization occurs, showing the $\log_2$-transformed fold change between the long-term peak magnitude and the first peak. The dashed line at 1 indicates a doubling in the long-term peak magnitude compared to the first peak. (c) Magnitude of the first response peak, representing the initial maximal output of the system before sensitization effects become significant. (d) Number of peaks required for the system's response to enter an $\epsilon$-ball ($\epsilon = 10^{-3}$) around the expected steady-state peak magnitude, where subsequent fluctuations remain within this neighborhood. Parameters: $\beta = 2$, $\mu = 1$, $\lambda = 1$, $\rho = 5$.

Beyond these constraints, the parameter region supporting robust sensitization is narrowed further by the need for a physiologically realistic number of priming pulses. Although sensitization remains possible for some parameter combinations, it may require an impractically large number of stimuli (see Figure 6d).

### IV. COMBINING SENSITIZATION AND HABITUATION

#### A. Hybrid Response: Sensitization with a Positive Feedback Loop

As illustrated in the main text, a hybrid response can be produced by adding a single negative interaction to a sensitization motif that contains a positive feedback loop. This addition creates a circuit whose output first increases and then decreases in magnitude.

Figure 7a shows a genetic circuit similar to Figure 4a, with a change in the output's promoter which makes it repressed by $I$ (LacI), which serves both as memory for the sensitization pathway as well as the habituation pathway.

The equations governing the system are as follows

$$\begin{aligned}
\frac{dX}{dt} &= \mu\,\theta^{-}(R) - \lambda\,X, \\
\frac{dI}{dt} &= \alpha\,\theta^{+}(X\,x) - \gamma\,I, \\
\frac{dR}{dt} &= \mu\,\theta^{-}(I) - \lambda\,R, \\
\frac{dG}{dt} &= \beta\,\theta^{+}(X\,x)\,\theta_2^{-}(I) - \lambda\,G,
\end{aligned} \tag{16}$$

where $\alpha$, $\beta$, $\mu$ are maximum promoter expression rates, which can be independently tuned to shape the system's dynamics. The separation of timescales is enforced by the basal degradation rate $\gamma$ for the slow memory protein $I$ and the increased rate $\lambda$ for the all other proteins, with $\lambda \geq \gamma$.

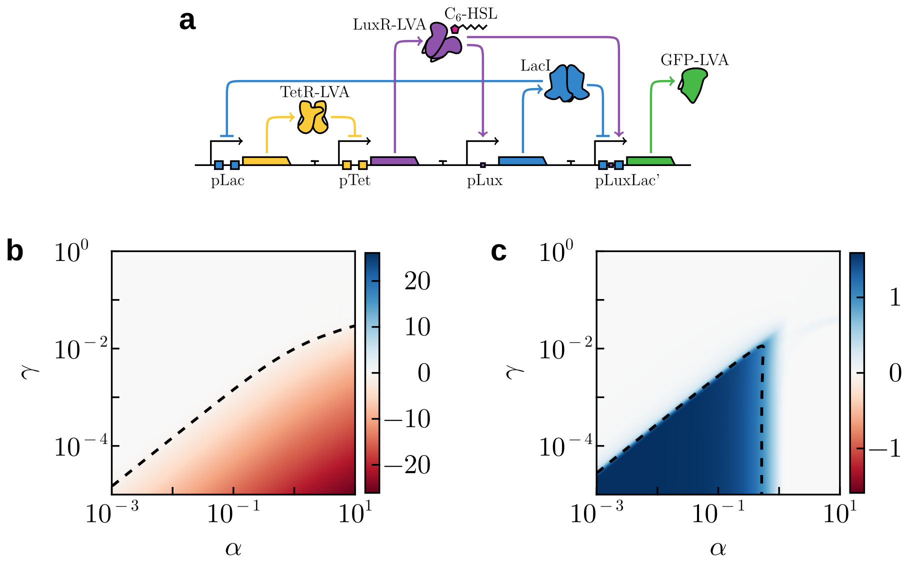


FIG. 7 **Hybrid Response: Sensitization with a Positive Feedback Loop.** (a) The proposed genetic circuit implementing the hybrid response by extending sensitization motif with a positive feedback loop Figure 4a. (b, c) Parameter spaces where habituation and sensitization occurs respectively, showing the $\log_2$-transformed fold change FC. The dashed line at 1 indicates doubling in the peak magnitude compared to the first peak, and dashed line at $-1$ indicates halving in the peak magnitude compared to the maximum peak in the trajectory. Parameters: $\beta = 4$, $\mu = 1$, $\lambda = 1$, $\rho = 1.5$.

Using a transfer function with a doubled half-activation constant for the repressor (I) concentration ($\theta_2^-(I)$) enhances the distinction between the two behaviors. This is a biologically plausible assumption, as a hybrid promoter's sensitivity would likely be hindered if not fine-tuned. In any case, this can effectively be optimized either through (directed) evolution, mutagenesis, or *in silico* models.

As shown in Figure 7b-c, the parameter ranges under which sensitization and habituation take place are similar to those of the individual circuits from which this extended motif is derived (Figure 1d, Figure 4d). Although these regions exhibit a partial overlap, the two behaviors cannot be optimized simultaneously. This is due to their divergent parameter dependencies: while both require a low memory degradation rate ($\gamma$), habituation monotonically strengthens with an increased maximal expression rate ($\alpha$), whereas sensitization weakens beyond an optimal $\alpha$ value, as it happens in the base sensitization motif. This inherent trade-off is a direct consequence of both behaviors sharing core circuit components for both behaviors.

#### B. Hybrid Response: Sensitization with Self-activation Loop

Similar to the previous hybrid circuit, the sensitization motif using a self-activation loop can also be extended in the same manner by replacing the output's promoter to be a hybrid promoter which can also be repressed, similar to the previous case (Figure 8a).

The equations governing the system are as follows:

$$\frac{dX}{dt} = \alpha\,\left[\varepsilon + (1-\varepsilon)\,\theta^+(X\,x)\right] - \gamma\,X, \tag{17}$$

$$\frac{dI}{dt} = \mu\,\left[\varepsilon + (1-\varepsilon)\,\theta^+(X\,x)\right] - \gamma\,I, \tag{18}$$

$$\frac{dG}{dt} = \beta\,\left[\varepsilon + (1-\varepsilon)\,\theta^+(X\,x)\right]\,\theta^-(I) - \lambda\,G, \tag{19}$$

where $\alpha$, $\mu$, and $\beta$ are maximum promoter expression, which can be independently tuned to shape the system's dynamics. Basal degradation rate $\gamma$ for untagged proteins, and increased rate $\lambda$ for tagged output protein, with $\lambda \geq \gamma$.

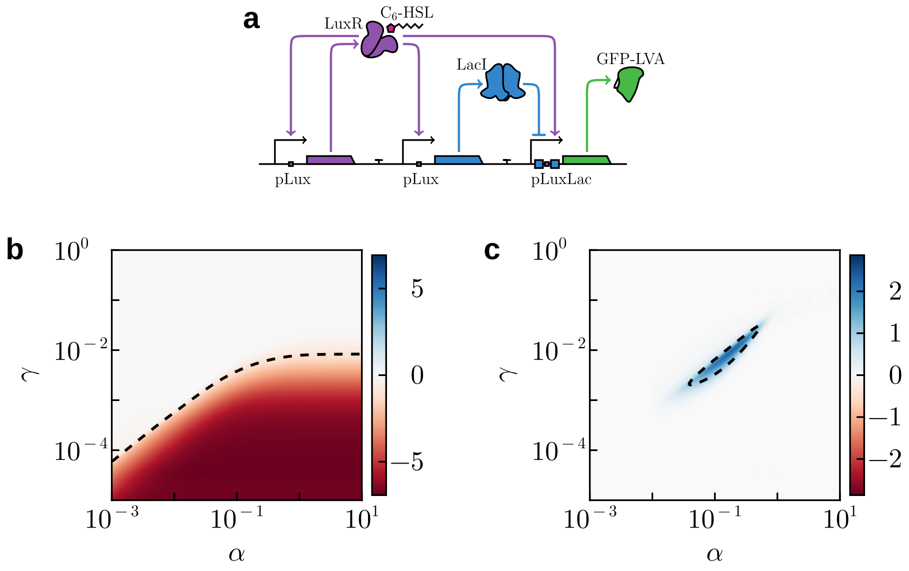


FIG. 8 **Hybrid Response: Sensitization with Self-activation Loop.** (a) The proposed genetic circuit implementing the hybrid response by extending sensitization motif with self-activation loop Figure 5a. (b, c) Parameter spaces where habituation and sensitization occurs respectively, showing the $\log_2$-transformed fold change FC. The dashed line at 1 indicates doubling in the peak magnitude compared to the first peak, and dashed line at $-1$ indicates halving in the peak magnitude compared to the maximum peak in the trajectory. Parameters: $\beta = 2$, $\mu = 10^{-1}$, $\lambda = 1$, $\varepsilon = 10^{-2}$.

As illustrated in Figure 8b-c, although habituation remains feasible, sensitization is significantly constrained by the requirement for two distinct, interdependent memory components. The parameter space in which sensitization occurs is narrower than the base sensitization case (Figure 5b). A further drawback is that the region supporting sensitization is disjoint from the region where habituation occurs.

Both receptor $X$ and repressor $I$ must persist over subsequent pulses to enable their accumulation over time. A significant challenge arises because maximum output expression coincides with the maximal expression of the repressor, as both proteins are under the positive regulation of the $X$–$x$ complex.

This constraint may be partially mitigated through further optimization of system parameters or by reducing the sensitivity of the repressor's promoter to the receptor complex. Nonetheless, the inherently limited parameter space in which the core sensitization motif works as intended would likely pose significant challenges for experimental implementation.

#### C. Hybrid Response: Sensitization with a Coherent Feed-Forward Loop

As in the case of sensitization motifs, an alternative hybrid response can also be created by extending the CFF sensitization circuit from Figure 6a.

The hybrid circuit (Figure 9a) can be implemented using several concurrent pathways. Receptor $X$ (LuxR-LVA) is constitutively expressed and binds the external input molecule $x$ upon its presence, forming the activator complex $X$–$x$, which activates downstream promoters. The output $G$ (GFP-LVA) is expressed from a hybrid promoter that is positively regulated by the $X$–$x$ complex and negatively regulated by repressor $I$.

Here, $I$ represents the total concentration of two variants of the same protein—one tagged and one untagged. Each variant is produced by one of two parallel pathways, which independently contribute to the overall behavior:

- A habituation pathway (Figure 1a), in which an untagged repressor $I_{\mathrm{H}}$ (LacI) is produced under direct activation by the $X$–$x$ complex.
- A sensitization pathway (Figure 6a), in which an additional repressor $R$ (TetR) is produced under direct activation by the $X$–$x$ complex. This repressor $R$, in turn, inhibits the expression of a tagged repressor $I_{\mathrm{S}}$ (LacI-LVA).

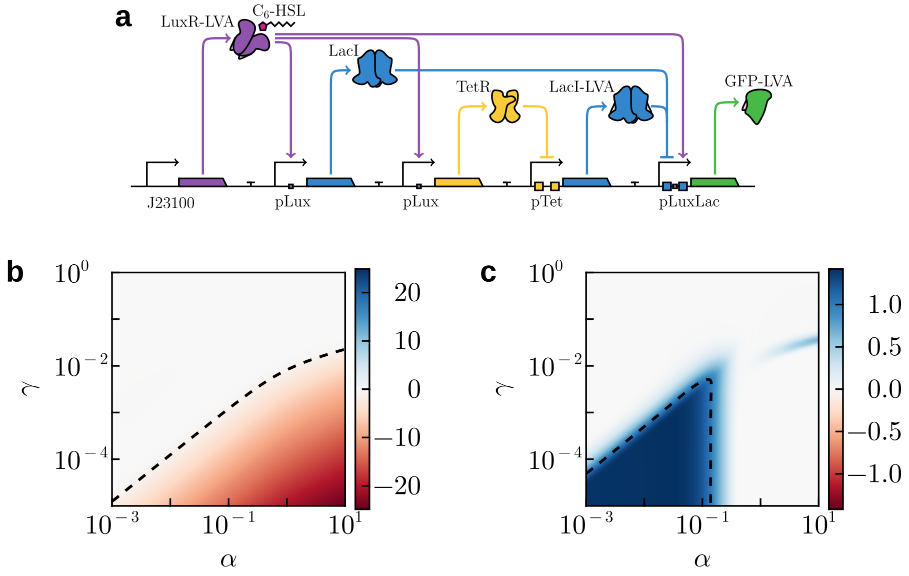


FIG. 9 **Hybrid Response: Sensitization with a Coherent Feed-Forward Loop.** (a) The proposed genetic circuit implementing the hybrid response by extending sensitization motif with a coherent feedforward loop Figure 6a. (b, c) Parameter spaces where habituation and sensitization occurs respectively, showing the $\log_2$-transformed fold change FC. The dashed line at 1 indicates doubling in the peak magnitude compared to the first peak, and dashed line at $-1$ indicates halving in the peak magnitude compared to the maximum peak in the trajectory. Parameters: $\beta = 2$, $\mu = 1$, $\lambda = 1$, $\rho = 5$.

This chain of repressors create a positive effect.

The total repressor concentration $I := I_{\mathrm{H}} + I_{\mathrm{S}}$ subsequently represses the output promoter.

In the absence of external input, the tagged repressor $I_{\mathrm{S}}$ is expressed at maximal levels, suppressing output production and impeding activation even when the activator complex is present. However, the $X$–$x$ complex can freely interact with the promoters regulating $I_{\mathrm{H}}$ and $R$, both of which exhibit similar dynamics and accumulate gradually over successive input pulses.

As the concentration of repressor $R$ increases, the tagged repressor $I_{\mathrm{S}}$ rapidly reaches a quasi-steady-state governed by the instantaneous level of $R$. Concentration of $I_{\mathrm{S}}$ diminishes with each successive pulse, thereby enabling progressively stronger activation of the output by the $X$–$x$ complex. Eventually, although $I_{\mathrm{S}}$ continues to decline, the untagged repressor $I_{\mathrm{H}}$ continues to accumulate independently, ultimately leading to renewed repression of output expression.

The equations governing the system are as follows

$$\frac{dX}{dt} = \mu - \lambda X, \tag{20}$$

$$\frac{dR}{dt} = \alpha\, \theta^{+}(X\, x) - \gamma\, R, \tag{21}$$

$$\frac{dI_H}{dt} = \alpha\, \theta^{+}(X\, x) - \gamma\, I_H, \tag{22}$$

$$\frac{dI_S}{dt} = \rho\, \theta^{-}(R) - \lambda\, I_S, \tag{23}$$

$$\frac{dG}{dt} = \beta\, \theta^{+}(X\, x)\, \theta^{-}(I_H + I_S) - \lambda\, G, \tag{24}$$

where $\alpha$, $\beta$, $\mu$, $\rho$ are maximum promoter expression rates, which can be independently tuned to shape the system's dynamics. The separation of timescales is enforced by the basal degradation rate $\gamma$ for the slow memory proteins and the increased rate $\lambda$ for the all other proteins, with $\lambda \geq \gamma$.

As shown in Figure 9b-c, the parameter ranges under which sensitization and habituation take place are similar to those of the individual circuits from which this extended motif is derived (Figure 1d, Figure 6d). Although these regions exhibit a partial overlap, the two behaviors cannot be optimized simultaneously. This is due to their divergent parameter dependencies: while both require a low memory degradation rate ($\gamma$), habituation monotonically strengthens with an increased maximal expression rate ($\alpha$), whereas sensitization weakens beyond an optimal $\alpha$ value, as it happens in the base sensitization motif.

This behavior resembles that of the first hybrid circuit presented (IV.A). In this case, however, there is no need to adjust promoter sensitivities; instead, distinct variants of the same protein with differing degradation dynamics are used.

Both implementations involve trade-offs. Although the use of a CFF increases the number of required components and thus the complexity, it also affords greater flexibility. Because the two behaviors are not coupled to the same memory component and instead rely on two separate proteins, their expression and degradation rates can be independently tuned, changing the point at which the circuit's behavior switches.

The same modularity could, in principle, be introduced into the positive feedforward loop case through the addition of components with distinct dynamics. However, when experimentally implementing such circuits, excessive flexibility and a large number of simultaneous optimization parameters can become a drawback. This is the primary reason for selecting the positive feedforward loop implementation in the main text, rather than the CFF-based design, even though both versions exhibit broadly similar behavior.

### V. MASSED–SPACED ACCUMULATION

A simplification for the massed–spaced regime can be made to facilitate the interpretation of results. Assume that all input pulses fully saturate the receptor's Hill function, and the output protein is expressed under a Heaviside step function $\mathcal{H}(\cdot)$ with the threshold set at the normalized half-saturation constant $K_{1/2} = 1$.

Under this assumption, the intermediate memory element $A$ is stimulated by $N$ instantaneous pulses of total strength $\alpha\Delta\tau_{\text{on}}$, spaced by a relaxation period of duration $\Delta\tau_{\text{off}}$, with the restriction that $\Delta\tau_{\text{on}} \ll \Delta\tau_{\text{off}}$. Its dynamics are governed by the equation:

$$\frac{dA}{dt} = \frac{\alpha\,\Delta\tau_{\text{on}}}{N} \sum_{k=0}^{N-1} \delta(t - t_k) - \gamma\, A, \tag{25}$$

where $t_k = k\,\Delta\tau_{\text{off}}$, and $\delta(t - t_k)$ denotes an impulsive input at time $t_k$. This formulation mirrors the structure of the memory accumulation equation (7), leading to qualitatively similar dynamics. In this case, however, the concern is not with the fine-grained dynamics of memory, but rather solely with its value at the start of the interval and whether it exceeds the threshold defined by the step transfer function.

The solution for $A(t)$ can be computed piecewise, starting from the initial condition $A(0) = 0$, since no input is initially present:

$$A(t) = \begin{cases} A(t_k^+)\, e^{-\gamma\,(t - t_k)} & \text{for } t_k < t < t_{k+1}, \\ A(t_k^-) + \dfrac{\alpha\,\Delta\tau_{\text{on}}}{N} & \text{at } t = t_k. \end{cases} \tag{26}$$

Letting $A_k \equiv A(t_k^+)$, the recurrence relation becomes:

$$A_k = A_{k-1}\, e^{-\gamma\,\Delta\tau_{\text{off}}} + \frac{\alpha\,\Delta\tau_{\text{on}}}{N},$$

with solution:

$$A_k = \frac{\alpha\,\Delta\tau_{\text{on}}}{N}\, \frac{1 - e^{-(k+1)\,\gamma\,\Delta\tau_{\text{off}}}}{1 - e^{-\gamma\,\Delta\tau_{\text{off}}}}, \tag{27}$$

which describes how much $A(t)$ has built up by the start of the $k$-th interval.

At long times after the last pulse, $t \gg t_{N-1}$, the memory decays freely:

$$\lim_{t\to\infty} A(t) = A_{N-1}\, e^{-\gamma\,(t-t_{N-1})} \approx 0,$$

thus the interest is in transient dynamics as the steady state will always be a complete lack of output.

Given the assumption of a step-like transfer function for the output, $G$ is produced at a constant rate $\beta$ when $A(t) \geq 1$, and decays otherwise. This leads to the differential equation:

$$\frac{dG}{dt} = \beta\,\mathcal{H}(A(t)-1) - \gamma\,G. \tag{28}$$

The duration of output production after the $k$-th input pulse is determined by the value of $A_k$, the state of $A(t)$ at the beginning of the interval after the instantaneous jump. Such crossing time is found by solving the equation governing the exponential decay of $A(t)$ from its initial value $A_k$:

$$A(t) = A_k\, e^{-\gamma\, t}.$$

Setting the desired output $A(t_k^*) = 1$, the time $t_k^* > t_k$ at which $A(t)$ decays to the threshold can be found by solving:

$$A(t_k^*) = 1 = A_k\, e^{-\gamma\,(t_k^*-t_k)} \implies t_k^* = t_k + \gamma^{-1}\,\log(A_k). \tag{29}$$

This gives the time it takes for $A(t)$ to decay from $A_k$ to the threshold where no more output will be produced, assuming no additional pulses are applied.

Since another pulse can arrive before the intermediate variable $A(t)$ decays below the threshold, the previous expression must be adjusted to account for such possibility. Accordingly, the duration of production during the $k$-th interval, denoted $\ell_k$, is defined as the length of time during which $A(t) \geq 1$:

$$\ell_k := \begin{cases} \min\left(t_k^* - t_k,\ t_{k+1} - t_k\right) & \text{if } A_k \geq 1 \text{ and } k < N-1, \\ t_k^* - t_k & \text{if } A_k \geq 1 \text{ and } k = N-1, \\ 0 & \text{otherwise.} \end{cases} \tag{30}$$

The last case arises when $A_k < 1$, in which case $t_k^*$ falls outside the $k$-th interval and no production occurs during that period.

This extension ensures that, for all intervals except the last, the production duration cannot exceed the time until the next pulse arrives. For the final interval, no subsequent pulse is applied, and hence the production duration is unbounded and extends until $A(t)$ naturally decays below the threshold.

The temporal evolution of output $G(t)$, equation (28), is thus a linear first-order ordinary differential equation subject to a piecewise constant forcing term. The analytical solution is thus contingent upon two distinct regimes, which depends upon the state of the memory $A(t)$.

When production is active—i.e., during periods where $A(t) \geq 1$—the Heaviside step function $\mathcal{H}(A(t)-1)$ equals 1. In this regime, the dynamics of $G(t)$ are governed by:

$$\frac{dG}{dt} = \beta - \gamma\,G \implies G(t) = G(t_0)\,e^{-\gamma\,(t-t_0)} + \frac{\beta}{\gamma}\left(1 - e^{-\gamma\,(t-t_0)}\right),$$

where $G(t_0)$ is the value of $G$ at the beginning of the interval.

Since $G$ is an independent observable that does not influence other system variables, the response to each input pulse can be treated independently. Focusing only on the contribution from the $k$-th pulse and ignoring residual buildup prior to $t_k$, the output during the active production phase is:

$$P_k(t) := \frac{\beta}{\gamma}\left(1 - e^{-\gamma\,(t-t_k)}\right), \qquad t \in [t_k, t_k + \ell_k).$$

Conversely, when production is inactive, i.e., $A(t) < 1$, the Heaviside function $\mathcal{H}(A(t)-1)$ vanishes and production is halted. The equation then reduces to the homogeneous form:

$$\frac{dG}{dt} = -\gamma\, G \implies G(t) = G(t_0)\, e^{-\gamma\,(t-t_0)},$$

describing pure exponential decay from its value at the end of the preceding production interval. Specifically, after the production phase from the $k$-th pulse halts, the produced output evolves as:

$$D_k(t) := \frac{\beta}{\gamma}\, \left(1 - e^{-\gamma\,\ell_k}\right) e^{-\gamma\,(t-t_k-\ell_k)}, \qquad t \geq t_k + \ell_k.$$

To obtain the complete expression $G(t)$ in systems characterized by finite input pulses, the net contributions from all previous pulses must be integrated, taking into account continuous degradation:

$$G(t) = \sum_{k=0}^{N-1} G_k(t), \qquad G_k(t) = \begin{cases} \frac{\beta}{\gamma}\, \left(1 - e^{-\gamma\,(t-t_k)}\right) & \text{if } t_k \leq t < t_k + \ell_k \\ \frac{\beta}{\gamma}\, \left(1 - e^{-\gamma\,\ell_k}\right) e^{-\gamma(t-t_k-\ell_k)} & \text{if } t \geq t_k + \ell_k \\ 0 & \text{otherwise,} \end{cases}$$

which can also be expressed using Heaviside functions as gatekeepers:

$$G(t) = \frac{\beta}{\gamma} \sum_{k=0}^{N-1} \left[ \left(1 - e^{-\gamma(t-t_k)}\right) \mathcal{T}_k + \left(1 - e^{-\gamma\,\ell_k}\right) e^{-\gamma\,(t-t_k-\ell_k)}\, \mathcal{H}(t - t_k - \ell_k) \right]$$

where $\mathcal{T}_k = \mathcal{H}(t-t_k) - \mathcal{H}(t-t_k-\ell_k)$ is a "boxcar" function created by a linear combination of step functions which is 1 only in the interval $t_k \leq t < t_k + \ell_k$ and 0 otherwise. Thus, $\mathcal{T}_k$ effectively isolates the sub-interval corresponding to the duration of the $k$-th production pulse. The step $\mathcal{H}(t - t_k - \ell_k)$ causes the output accumulated from that pulse to decay exponentially towards zero, continuing after the pulse has ended.

The expression can be further compacted by defining the following two functions:

- $\tau_k^+(t) := \min(t - t_k,\ \ell_k)$, representing the duration of output production in the $k$-th interval up to time $t$, bounded by the interval duration
- $\tau_k^-(t) := \max(0,\ t - t_k - \ell_k)$, representing the time elapsed since production halted in the $k$-th interval, bounded by the time since the interval ended if production continued beyond its duration.

$$G(t) = \frac{\beta}{\gamma} \sum_{k=0}^{N-1} \left(1 - e^{-\gamma\,\tau_k^+(t)}\right) e^{-\gamma\,\tau_k^-(t)}\, \mathcal{H}(t - t_k).$$

Assuming that during the active production phase, the production rate $\beta$ exceeds the decay rate $\gamma$ ($\beta > \gamma$), the peak output for the $k$-th interval is reached at time $t_k + \ell_k$, which marks the precise moment when active production stops, followed by an exponential decay of $G(t)$. Considering that all production intervals are equally spaced, and the lowest possible starting output ($G_k = 0$) occurs in the very first interval $k = 0$, it logically follows that every subsequent interval will achieve a peak output at least as high as the previous one.

Therefore, the absolute maximum output over the entire trajectory will be the last output peak, $G(t_{N-1}+\ell_{N-1})$.

#### A. Simple Example of Massed versus Spaced

To simplify the derivation of the conditions under which spaced learning is more effective than massed learning, the following development will be under the assumption that

$$\log\left(\frac{\alpha\,\Delta\tau_{\text{on}}}{N}\right) \geq \gamma\,\Delta\tau_{\text{off}}, \tag{31}$$

which ensures that when there are multiple pulses, the next pulse arrives before the production halts. There may exist an extended parameter regime in which spaced learning can outperform massed learning; however, analytical tractability of this regime requires a more rigorous treatment (see the extended space in Figure 10b).

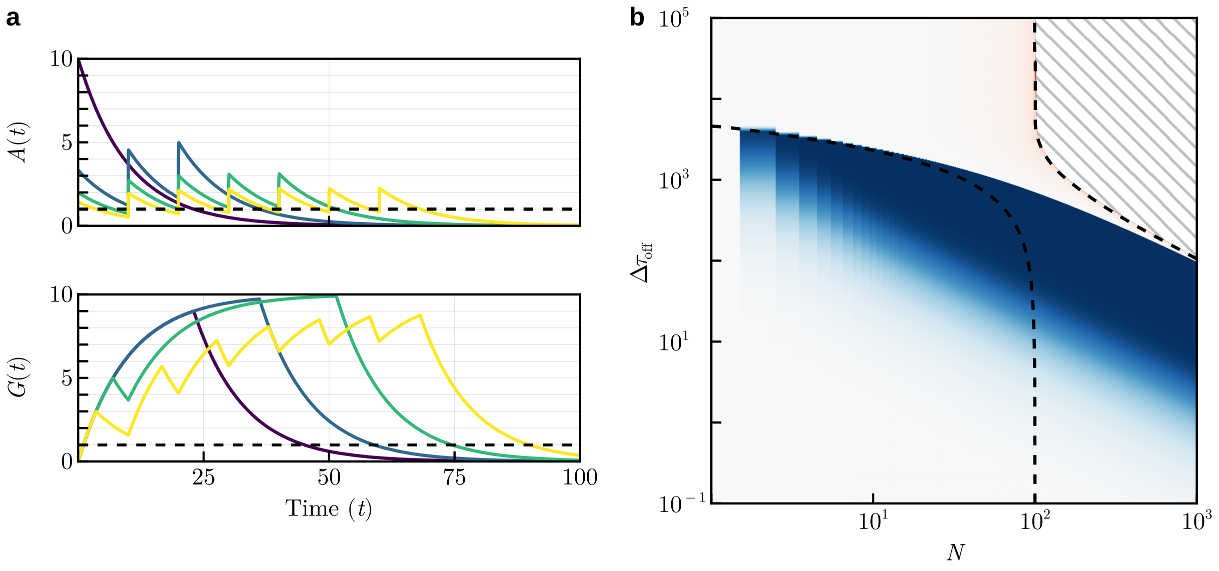


FIG. 10 In (a), a sample of timeseries for the system is shown, with $N$ instant inputs applied with a relaxation period in between. The massed case ($N = 1$, ■) reaches the maximum peak faster, but the peak height can be enhanced by dividing the input. It is exemplified with $N = 3$ pulses (■), and reaching its maximum when $N = 5$ (■), which is a case outside the assumption (31) as output production halts before the first relaxation phase ends. Further dividing the input ($N = 7$, ■) can be detrimental, even though the time output $G$ is above a normalized threshold continues increasing. Parameters: $\alpha = 10^2$, $\beta = 1$, $\gamma = 10^{-1}$, $\Delta\tau_{\rm on} = 10^{-1}$, $\Delta\tau_{\rm off} = 10^1$. In (b), the parameter space, divided into three distinct regimes: (i) A regime for small number of pulses $N$ and short relaxation periods $\Delta\tau_{\rm off}$, equation (31) holds, leading to continuous production that persists until after the last pulse; (ii) An intermediate regime where production halts during portions of the relaxation phases, yet repeated stimulation can accumulate memory across pulses and enhance the maximum output produced; (iii) A regime with a large number of pulses and long relaxation times, where the input is diluted and fragmented to build up enough memory for any output to be generated, filled with a gray hatching. Parameters: $\alpha = 10^5$, $\beta = 1$, $\gamma = 10^{-3}$, $\Delta\tau_{\rm on} = 10^{-3}$.

#### Maximum Peak for $N = 1$

For a single pulse ($N = 1$), output production begins immediately after the input pulse and continues until the memory element A(t) decays below 1. The duration of this production, $\ell_0$, is given by:

$$\ell_0 = \gamma^{-1} \log(A_0) = \gamma^{-1} \log(\alpha\, \Delta\tau_{\rm on}),$$

at which point the output reaches its maximum

$$\max_t G(t)|_{N=1} = G(\ell_0) = \frac{\beta}{\gamma} \left(1 - e^{-\gamma\, \ell_0}\right) = \frac{\beta}{\gamma} \left(1 - \frac{1}{\alpha\, \Delta\tau_{\rm on}}\right), \tag{32}$$

given that $e^{-\gamma\, \ell_0} = e^{-\log(\alpha\, \Delta\tau_{\rm on})} = (\alpha\, \Delta\tau_{\rm on})^{-1}$.

#### Maximum Peak for $N = 2$

For the case where two pulses of half duration are applied ($N = 2$), with a relaxation period of length $\Delta\tau_{\rm off}$, a first pulse will be applied as before. Due to the assumption (31), output will be produced during the whole relaxation period ($\ell_0 = \Delta\tau_{\rm off}$). After the second input, production will continue until the memory element A(t) decays below 1. Given $A$ at the start of the second pulse following equation (27) is

$$A_1 = \frac{\alpha\, \Delta\tau_{\rm on}}{2} \, \frac{1 - e^{-2\,\gamma\, \Delta\tau_{\rm off}}}{1 - e^{-\gamma\, \Delta\tau_{\rm off}}} = \frac{\alpha\, \Delta\tau_{\rm on}}{2} \left(1 + e^{-\gamma\, \Delta\tau_{\rm off}}\right)$$

the duration of the production phase is given by:

$$\ell_1 = \gamma^{-1} \log(A_1) = \gamma^{-1} \log\left(\frac{\alpha\, \Delta\tau_{\rm on}}{2} \left(1 + e^{-\gamma\, \Delta\tau_{\rm off}}\right)\right).$$

In this case the absolute maximum output $G$ will be reached at $t = \Delta\tau_{\text{off}} + \ell_1$, the point where no more output will be produced:

$$G(\Delta\tau_{\text{off}} + \ell_1) = \frac{\beta}{\gamma}\left(\left(1 - e^{-\gamma\,\Delta\tau_{\text{off}}}\right)\, e^{-\gamma\,\ell_1} + 1 - e^{-\gamma\,\ell_1}\right) = \frac{\beta}{\gamma}\left(1 - e^{-\gamma\,(\Delta\tau_{\text{off}}+\ell_1)}\right),$$

which can be further simplified by substituting $\ell_1$:

$$\max_t G(t)|_{N=2} = G(t_1^*) = \frac{\beta}{\gamma}\left(1 - \frac{2\,e^{-\gamma\,\Delta\tau_{\text{off}}}}{\alpha\,\Delta\tau_{\text{on}}\,(1 + e^{-\gamma\,\Delta\tau_{\text{off}}})}\right), \qquad (33)$$

with $t_1^* = \Delta\tau_{\text{off}} + \ell_1$ the time it takes for output production to stop.

The condition for a two-pulse spaced input to produce a higher peak response than its massed counterpart is determined by solving

$$\max_t G(t)|_{N=2} > \max_t G(t)|_{N=1}$$

which simplifies algebraically to the condition:

$$\frac{\beta}{\gamma}\left(1 - \frac{2e^{-\gamma\Delta\tau_{\text{off}}}}{\alpha\Delta\tau_{\text{on}}(1 + e^{-\gamma\Delta\tau_{\text{off}}})}\right) > \frac{\beta}{\gamma}\left(1 - \frac{1}{\alpha\Delta\tau_{\text{on}}}\right).$$

Assuming all parameters are positive, this inequality simplifies to:

$$e^{-\gamma\,\Delta\tau_{\text{off}}} < 1,$$

which holds strictly whenever $\gamma\,\Delta\tau_{\text{off}} > 0$. Since both the decay rate $\gamma$ and the relaxation period length $\Delta\tau_{\text{off}}$ are positive, this condition is always satisfied. Thus, the spaced protocol consistently yields a higher peak response than the massed one under the previous assumption (31).

In the limit of no degradation, $\gamma \to 0^+$, memory decay becomes negligible, causing the accumulated signal to persist indefinitely. As a result, output production effectively stays "on" continuously, and any temporal separation between inputs provides no additional benefit. Similarly, when spacing between pulses banishes, $\Delta\tau_{\text{off}} \to 0^+$, the pulses are delivered almost simultaneously, converging to a single massed input.

#### Maximum Peak for $N > 1$

Following a similar process, the previous condition can be generalized for the maximum peak given $N > 1$ pulses, which is the value of output $G$ evaluated at $t_N = (N-1)\,\Delta\tau_{\text{off}} + \ell_{N-1}$:

$$G(t_{N-1}^*) = \frac{\beta}{\gamma}\left(1 - \frac{N\,e^{-\gamma\,(N-1)\,\Delta\tau_{\text{off}}}}{\alpha\,\Delta\tau_{\text{on}}\,S_N}\right),$$

where $S_N = 1 + e^{-\gamma\,\Delta\tau_{\text{off}}} + \cdots + e^{-(N-1)\gamma\Delta\tau_{\text{off}}}$ is a sum of a finite geometric series. This can be reduced to

$$\max_t G(t)|_N = G(t_{N-1}^*) = \frac{\beta}{\gamma}\left(1 - \frac{Ne^{-\gamma(N-1)\,\Delta\tau_{\text{off}}}}{\alpha\,\Delta\tau_{\text{on}}}\,\frac{1 - e^{-\gamma\,\Delta\tau_{\text{off}}}}{1 - e^{-N\,\gamma\,\Delta\tau_{\text{off}}}}\right) \qquad (34)$$

The condition

$$\max_t G(t)|_N > \max_t G(t)|_{N=1}$$

simplifies algebraically to:

$$N\,e^{-\gamma\,(N-1)\,\Delta\tau_{\text{off}}}\,\frac{1 - e^{-\gamma\,\Delta\tau_{\text{off}}}}{1 - e^{-N\,\gamma\,\Delta\tau_{\text{off}}}} < 1,$$

which holds true for all $N > 1$ and $\gamma\,\Delta\tau_{\text{off}} > 0$.

To prove this inequality, consider the following:

Let $x = e^{-\gamma\,\Delta\tau_{\text{off}}}$, so that $0 < x < 1$. Substituting into the original expression yields:

$$N\,x^{N-1}\,\frac{1-x}{1-x^N} < 1.$$

Using the factorization $1 - x^N = (1-x)\sum_{k=0}^{N-1} x^k$, the inequality becomes:

$$\frac{N\,x^{N-1}}{\sum_{k=0}^{N-1} x^k} < 1,$$

Since $\sum_{k=0}^{N-1} x^k$ is positive the following are equivalent inequalities:

$$N\,x^{N-1} < \sum_{k=0}^{N-1} x^k \iff \sum_{k=0}^{N-1} x^k - N\,x^{N-1} > 0.$$

Define the function:

$$S(x) = \sum_{k=0}^{N-1} x^k - Nx^{N-1},$$

and analyze its limits:

- As $x \to 0^+$, each term $x^k \to 0$ except for $x^0 = 1$, so $\lim_{x\to 0^+} S(x) \to 1$.
- If $x = 1$, $S(1) = N - N = 0$.

Moreover, since $0 < x < 1$, the sequence $\{x^k\}_{k=0}^{N-1}$ is strictly decreasing. Therefore, the arithmetic mean of these terms satisfies:

$$\frac{1}{N}\sum_{k=0}^{N-1} x^k > x^{N-1} \iff \sum_{k=0}^{N-1} x^k > Nx^{N-1},$$

which is precisely $S(x) > 0$.

Thus, the inequality holds for all integers $N > 1$ given $\gamma\,\Delta\tau_{\text{off}} > 0$.
